# *E. coli* RsmF activity depends on prior modification of 16S rRNA helix 44

**DOI:** 10.64898/2026.07.05.736617

**Authors:** Mohamed I. Barmada, Aya Hanna, Camden R. Bair, Erin N. McGinity, Natalia Zelinskaya, Debayan Dey, Graeme L. Conn

**Affiliations:** Department of Biochemistry, Emory University School of Medicine, Atlanta, Georgia, 30322, USA; Graduate Program in Biochemistry, Cell and Developmental Biology (BCDB), Emory University, Atlanta, Georgia, 30322, USA; Graduate Program in Microbiology and Molecular Genetics (MMG), Emory University, Atlanta, Georgia, 30322, USA; Department of Microbiology and Immunology, Emory University School of Medicine, Atlanta, Georgia, 30322, USA

**Author notes:** Co-first authors (Mohamed I. Barmada, Aya Hanna).

**Keywords:** 16S rRNA, RNA modification, methyltransferase, SHAPE-MaP

## Abstract

Bacterial ribosomal RNA (rRNA) methylations are important for accurate translation. Four distinct methylations incorporated by RsmE, RsmF, and RsmH/ RsmI form a cluster of three modified 16S rRNA nucleotides (m^3^U1498, m^5^C1407, and m^4^Cm1402) surrounding the decoding center of the 30S subunit. Given their common substrate requirement of a late-stage intermediate 30S subunit, these enzymes likely act contemporaneously during subunit biogenesis, but whether there exists a required modification order is unknown. Here, using hypomethylated 30S subunits obtained from a collection of *rsmH/I/E/F*-deleted *Escherichia coli* strains, we identify RsmF activity to be highly dependent on prior modification of h44 both *in vitro* and in *E. coli*. RsmF activity on hypomethylated 30S subunits could be partially rescued by prior *in vitro* methylation using RsmE and RsmH, indicating that incorporation of these methyl groups directly shapes h44 for recognition by RsmF. RNA structure probing using SHAPE-MaP and molecular dynamics simulations reveal specific alterations in 16S rRNA structure and dynamics in the absence of the m^4^C1402 (RsmH) and m^3^U1498 (RsmE) modifications that likely restrict RsmF action. These studies thus uncover a previously unappreciated “order of operations” for 16S rRNA modification during ribosome biogenesis with important implications for studies on the collective functions of these modifications.

**Graphical Abstract:** 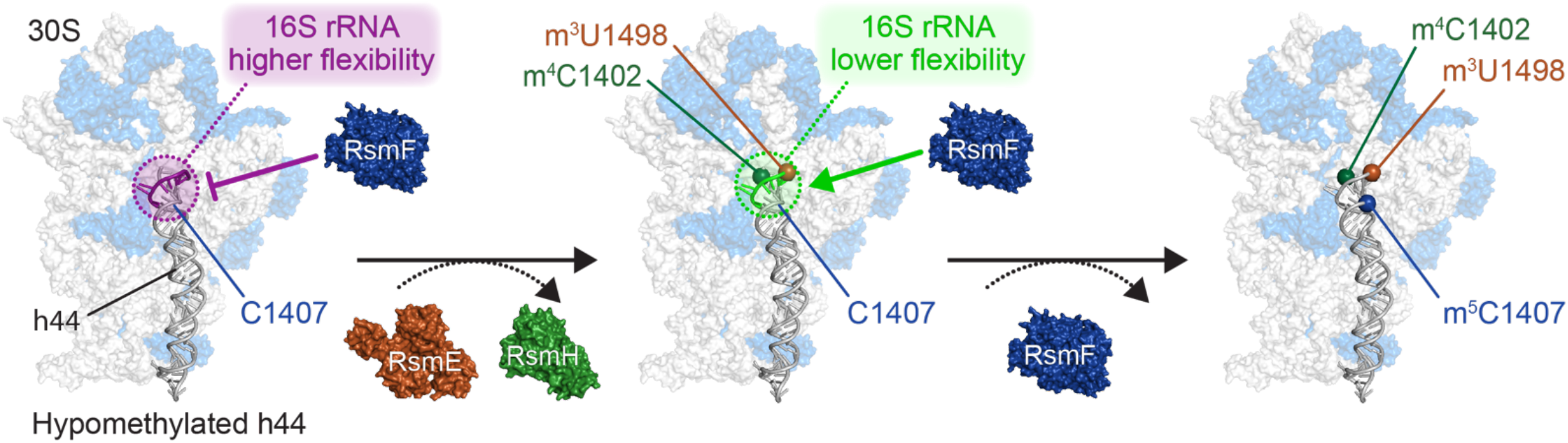

**Key Points:**

- 16S rRNA C1407 modification by RsmF depends on the prior action of RsmE and RsmH in E. coli
- m^5^C1402/ m^3^U1498 alter 16S rRNA nucleotide dynamics creating a 30S substrate suitable for RsmF
- An order of operations exists for h44 modifying enzymes acting at the 30S subunit decoding center

## INTRODUCTION

The bacterial ribosome undergoes extensive post-transcriptional modification through methylation of its RNA components. Predominantly clustered at the functional centers of both ribosomal subunits (1), methylated nucleotides play roles in subunit assembly (2,3), fine-tune translational fidelity (4-6), promote ribosome-ligand interactions (7), and ensure appropriate responses to stress (8,9). A prominent example of such clustering is the presence of four modifications on three 16S rRNA nucleotides, m^3^U1498, m^5^C1407, and m^4^Cm1402, surrounding the decoding center at the aminoacyl-(A) and the peptidyl-tRNA (P) sites on helix 44 (h44) of the small (30S) ribosomal subunit (**Fig. 1A**). Ubiquitously present across bacteria, these four methyl groups are deposited by the conserved S-adenosyl-L-methionine (SAM)-dependent methyltransferases RsmE (m^3^U1498), RsmF (m^5^C1407), RsmH (m^4^C1402), and RsmI (Cm1402) (5,10,11).

**Fig. 1.**
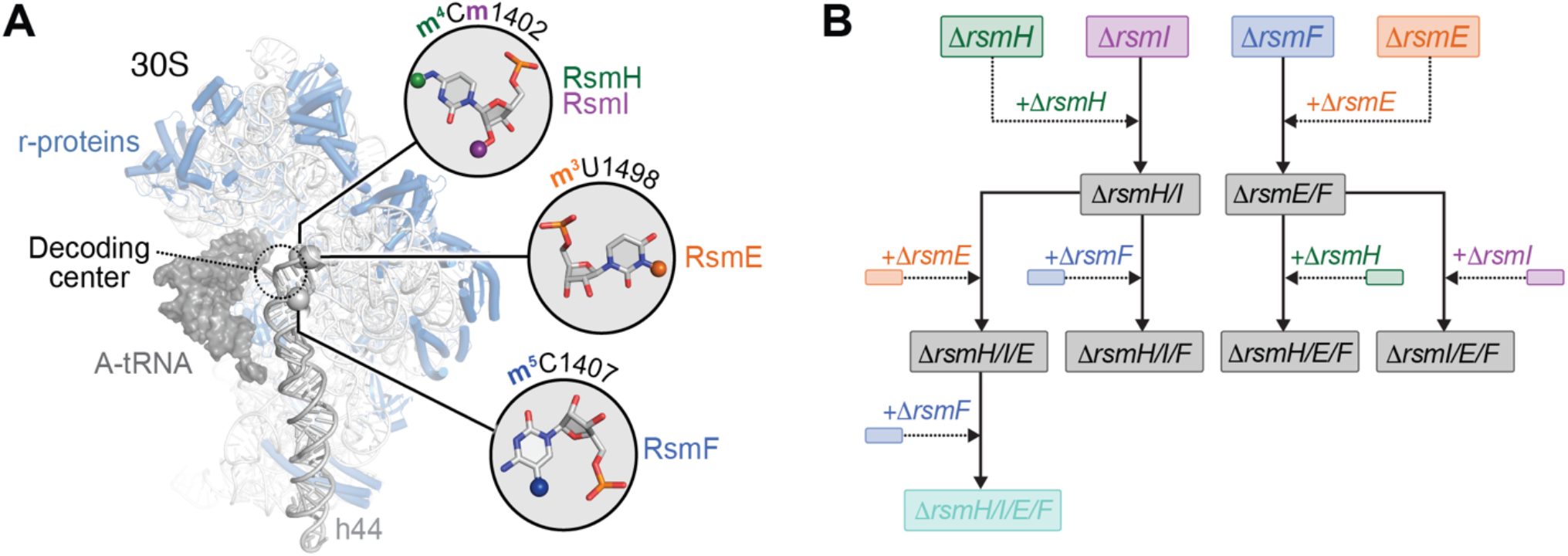
h44 is marked by a cluster of modifications surrounding the decoding center. (**A**) Structure of the *E. coli* 30S subunit (from PDB 8IFB) (39) with the cluster of three modified nucleotides highlighted and shown in zoomed-in views. The A-site tRNA (gray surface) is shown for reference adjacent to the decoding center and small subunit ribosomal proteins are shown in blue. (**B**) Workflow for generation of double, triple and quadruple Rsm deletion strains. Solid line arrows denote connections from recipient strain to new deletion and dashed line arrows indicate the DNA source for P1 phage transduction.

Although none of the modifications, nor the enzymes which incorporate them, are individually essential for bacterial growth, their loss results in various modest but discernable effects on bacterial fitness and ribosome function. For instance, deletion of either RsmI or RsmH (m^4^Cm1402) has been associated with translational fidelity defects and increases in bacterial susceptibility to various forms of stress (5,12-14). Likewise, loss of RsmF (m^5^C1407) increases ectopic protein expression (15) and alters the kinetics of proper codon-anticodon recognition (16), whereas absence of RsmE (m^3^U1498) causes increased resistance to antibiotics (17). Considering the proximity of these nucleotides, there is clear potential for these modifications to collectively shape the local environment around the decoding center. However, the combined influence of these modifications on the function of the A and P sites has not been directly explored.

Prior studies suggest that each h44 methyl group is likely incorporated late during 30S biogenesis with all of RsmE, RsmF, RsmH and RsmI only being able to methylate a mature 30S subunit, but not naked 16S rRNA, *in vitro* (5,11,18). Thus, with the likely substrate for these enzymes in cells being a late-stage assembly intermediate or a mature 30S subunit, the modifications not only occupy adjacent positions within h44, but are likely also deposited near-contemporaneously during 30S assembly. As such, the question arises of whether there exists any dependency among these enzymes RsmH, RsmI, RsmE, and RsmF on the prior action of one or more of the others. To investigate whether there is a potential hierarchy of methylation within h44–and as a platform for future studies to investigate how these modifications collectively shape 30S subunit assembly and function–we generated a series of *Escherichia coli rsm* gene deletion strains lacking combinations of two, three, or all four of the enzymes responsible for incorporating the h44 methyl modifications.

Using 30S subunits isolated from these deletions strain, we show that the m^5^C1407 methyltransferase RsmF depends on the prior action of other intrinsic 16S rRNA methyltransferases for its activity *in vitro* and in the bacterial cell. In particular, m^3^1498 and m^4^C1402, deposited by RsmE and RsmH, respectively, are sufficient to rescue RsmF activity on hypomethylated 30S subunits *in vitro*. RNA structure probing of 16S rRNA from hypomethylated 30S subunits that support a range of RsmF activities, reveal that the m^3^1498 and m^4^C1402 modifications alter h44 nucleotide dynamics opposite C1407, restricting 30S substrate recognition by RsmF. Collectively, these results reveal a hierarchy of modification within h44 and offer insights into the effects of rRNA methylation on the functions of ribosomal assembly factors during 30S biogenesis.

## MATERIAL AND METHODS

### Generation of E. coli 16S rRNA (h44) methyltransferase deletion strains

Starting from three single gene *E. coli* deletion strains Δ*rsmH*, Δ*rsmE*, and Δ*rsmF* from the Keio Collection (derived from *E. coli* BW25113 and with a kanamycin resistance marker; Kan^R^) (19), as well as a previously generated Δ*rsmI* strain (with a chloramphenicol resistance marker; Cam^R^) (20), new strains with multiple methyltransferase gene deletions were generated iteratively using P1 phage transduction (summarized in **Fig. 1B**) (21). Where required, the Kan^R^ marker was removed from recipient strains using the pCP20 plasmid, encoding FLP recombinase, and verified by PCR. The Δ*rsmH* and Δ*rsmE* genotypes were first transferred to the Δ*rsmI* and Δ*rsmF* backgrounds, respectively, to generate Δ*rsmH/I* and Δ*rsmE/F* double deletion strains. Next, Δ*rsmE* and Δ*rsmF* genotypes were transferred to the Δ*rsmH/I* background to generate the triple deletion strains Δ*rsmH/I/E* and Δ*rsmH/I/F*, and the Δ*rsmH* and Δ*rsmI* genotypes were transferred to the Δ*rsmE/F* background to generate the Δ*rsmH/E/F* and Δ*rsmI/E/F* strains. Lastly, the quadruple Δ*rsmH/I/E/F* strain was generated independently in two ways, by transferring the Δr*smF* genotype to the Δ*rsmH/I/E* background.

As noted above, the Kan^R^ marker was flipped out of new deletion strains to enable subsequent genetic transfers, except for Δ*rsmH/E/F* and Δ*rsmI/E/F* strains which were not used as recipients for additional deletions and thus retain kanamycin resistance. Additionally, Cam^R^ was maintained in all strains with Δ*rsmI*. Strain genotypes and Kan^R^ removal were validated by PCR (**Supplementary Table S1**) and whole genome sequencing (NCBI BioProject accession PRJNA1482076).

### Wild-type and hypomethylated 30S subunit isolation

30S subunits from *E. coli* wild-type BW25113 and deletion strains were isolated following the same general procedure with all centrifugation and dialysis steps performed at 4 °C. All strains were grown at 37 °C in lysogeny broth (LB) to mid-log phase (~0.6 absorbance at 600 nm) at which time the cells were harvested by centrifugation and washed twice with Buffer 1 and once with Buffer 2 (buffer compositions are provided in **Supplementary Table S2**). Next, cells were resuspended in Buffer 2 (9 ml/ g wet cell pellet) supplemented with TURBO DNase (1 unit/ ml lysate; Invitrogen) and lysed using an LM20 microfluidizer (Microfluidics). Cell debris was cleared by two successive centrifugation steps (17,000 and 27,000 × *g*), and the resulting supernatant spun at ~280,000 × *g* for 16 hours to pellet the ribosomal fractions. The pellets were resuspended in Buffer 2 before being layered in 100 OD_260_ (~500-700 μl) fractions on 10-30% sucrose gradients (38.5 ml total volume) made under low-Mg^2+^ using Buffer 3 (**Supplementary Table S2**). The sucrose gradients were centrifuged at ~95,000 × *g* for 16 hours, and the separated ribosomal subunits were fractionated using an ÄKTA Purifier 10 System. Fractions containing 30S subunits were pooled, pelleted (~220,000 × *g* for 18 hours), and resuspended in 300-500 μl Buffer 2. The resuspended subunits were dialyzed against Buffer 2 before being frozen for later use. Prior to use, 30S subunits were activated by incubation at 42 °C for 5 minutes and then room temperature for 10 minutes, similarly as to what was done before (22).

### Rsm protein expression and purification

Genes encoding the four intrinsic h44-modifying methyltransferases, RsmE, RsmF, RsmH, and RsmI, were PCR-amplified (**Supplementary Table S1**) from *E. coli* BW25113 genomic DNA and subcloned into a modified pET44a vector with an N-terminal hexahistidine (6×His) tag, creating plasmids pET44-RsmE, pET44-RsmF, pET44-RsmH, and pET44-RsmI. pET44-RsmI is the same plasmid as previously reported (20). Each vector was individually used to transform BL21(DE3) cells for IPTG-induced protein expression with subsequent purification accomplished using sequential Ni^2+^-affinity and size exclusion chromatographies essentially identically as described previously (20). The specific buffer composition differed slightly for each protein as detailed in **Supplementary Table S2**. The G194R-RsmE and the D55R-RsmH variants were generated using MEGAWHOP mutagenesis (23) (**Supplementary Table S1**) and purified using Ni^2+^-affinity spin columns (New England BioLabs). Protein folding/ stability and prep-to-prep quality control was done using nano differential scanning fluorimetry on a NanoTemper Tycho NT.6.

### [^3^H]-SAM *in vitro* methyltransferase assay

*In vitro* methyltransferase activity assays were conducted as described previously (20). Briefly, enzyme and 30S subunit were incubated with [^3^H]-SAM in Buffer G (10 μl total reaction volume; **Supplementary Table S2**) at 37 °C and reactions quenched with 90 μl of 5% ice-cold trichloroacetic acid (TCA). Quenched reactions were applied to a glass microfiber filter and free [^3^H]-SAM removed by washing 5% TCA (with 200 μl). The ^3^H remaining on the filter, as counts per minute (cpm; corresponding to 30S subunit methylation) was then determined by scintillation counting (Beckman Coulter LS6500). Data were plotted in Graphpad Prism 11 after subtraction of background values derived from corresponding control assay without enzyme.

Initial methylation assays comparing enzyme activity with 30S from wild-type and singly-deleted h44 methyltransferase *E. coli* strains were run for 60 minutes at an enzyme:30S substrate molar ratio of 1:1 for RsmE and RsmI (100 nM) and 4:1 for RsmH and RsmF (800 nM enzyme and 200 nM 30S subunit). Time course experiments were performed at similar enzyme:30S ratios, and the reaction mixtures were quenched at 0, 2.5, 5, 10, 20, 40, 60, and 90 minutes for RsmE, RsmF and RsmI, and at 0, 5, 15, 30, 60, and 90 minutes for RsmH. All additional single-time point experiments were performed for 30 minutes at the same enzyme:30S subunit ratios but using 200 nM RsmE or RsmI. Reactions indicated as “10×” relative enzyme concentration contained 10 times the above-mentioned enzyme concentrations (with substrate concentration remaining fixed). The [^3^H]-SAM concentration was kept constant at 0.84μM in all assays. For single-time point experiments where data is normalized, the condition/substrate with the highest values was chosen as the reference; values measured as lower than background are reported as zero for these experiments.

To prepare *in vitro* RsmE/RsmH methylated 30S subunits for the RsmF methylation rescue assays, 2 μM Δ*rsmH/E/F*-, Δ*rsmH/I/E/F*-, and Δ*rsmF*-derived 30S subunits were separately incubated with RsmE (3 μM), RsmH (5 μM), and non-radiolabeled SAM (20 μM) in Buffer G for 90 minutes at 37 °C. The reaction mixture was then passed through a NEBExpress Ni^2+^-Spin Column to separate the methylated 30S subunit (in the flowthrough) from the 6×His-tagged RsmE and RsmH enzymes (which remained bound to the column). Methylated 30S subunits were then dialyzed against Buffer 2 before use. For experiments to assess and account for substrate damage throughout this treatment, separate aliquots of the Δ*rsmH/I/E/F* and Δ*rsmF* 30S subunits underwent identical treatment but without RsmE and RsmH addition (mock). Assays with RsmE/RsmH-methylated 30S subunit and those with the catalytically inactive RsmE-G194R and RsmH-D55R were conducted as described above for the other single-time point experiments.

### 16S rRNA bisulfite sequencing

16S rRNA was extracted from purified wild-type, Δ*rsmF*, Δ*rsmH/I/E*, and Δ*rsmH/I/E/F* 30S subunits using a Monarch Spin RNA Cleanup kit (New England Biolabs). RNA bisulfite conversion and clean up was done using the Methylamp RNA Bisulfite Conversion Kit (EPIGENTEK). Bisulfite-treated 16S rRNA was reverse transcribed with SuperScript III (Illumina), and the resulting cDNA product desalted on a G25 spin column (Cytiva) and PCR amplified with Q5 Hot Start High-Fidelity DNA Polymerase (New England Biolabs) using bisulfite-specific primers for both steps, as previously described (24) For PCR, primers were used that amplify a ~100 nucleotide region encompassing C1407 (and C1402) and C967 (**Supplementary Table S1**). The resulting ~100 bp amplicons were purified using a Monarch PCR & DNA Cleanup Kit (New England Biolabs), blunt-end ligated into the pCR Blunt II-TOPO vector using the Zero Blunt TOPO PCR Cloning Kit (Invitrogen), and used to transform chemically competent *E. coli* DH5α cells. For each 16S rRNA, derived from wild-type, Δ*rsmF*, Δ*rsmH/I/E*, or Δ*rsmH/I/E/F* 30S subunits, sets of either five (for the “control” C967 amplicon) or 10 (for C1407 amplicon) single colonies were cultured overnight at 37 °C in LB medium and the individual plasmids isolated using a QIAprep Spin Miniprep Kit (QIAGEN) and for Sanger sequencing (GENEWIZ). An additional set of 10 individual colonies (processed identically) were examined to further verify the result for the Δ*rsmH/I/E* and C1407 amplicon combination. The extent of bisulfite conversion (expected C, sequenced as T) at the methylated nucleotides (C967, C1402, and C1407) was calculated by dividing the number of non-protected (C to T) base calls at the methylated nucleotide positions by the total number of sequenced plasmids.

### Selective 2′-hydroxyl acylation analyzed by primer extension and mutational profiling (SHAPE-MaP)

#### Library preparation

Purified 30S subunits were treated with either DMSO (control) or SHAPE reagent 2A3, prepared in-house as previously described (25), for 15 minutes at 37 °C. The reaction was stopped by addition of an equal volume of 1 M dithiothreitol and incubation for 5 minutes at room temperature. Two or three independent replicates were performed per 30S subunit type analyzed (as noted in the associated figures). 16S rRNA was extracted from the modified 30S subunits using a Monarch Spin RNA Cleanup kit (New England Biolabs) and reverse transcribed with Superscript II (Invitrogen) in MaP buffer following procedures established previously (26) (**Supplementary Tables S1** and **S2**). The resulting cDNA were desalted using G25 spin columns (Cytiva) and PCR amplified using Q5 Hot Start High-Fidelity DNA Polymerase (New England Biolabs). Primers contained partial Nextera XT Illumina sequencing adapters and inline barcodes and were designed to amplify the 16S rRNA region from C1367 to A1542 (encompassing h44 and h45; **Supplementary Table S1**). The resulting amplicons were cleaned up using a Monarch PCR & DNA Cleanup Kit (New England Biolabs), their purity was verified via gel electrophoresis, and concentrations determined using measurement on a Nanodrop 2000c Spectrophotometer (Thermo Scientific) and Qubit4 fluorometer (Invitrogen). Libraries were then constructed with the Nextera XT DNA library Preparation Kit (Illumina, FC-131-1096) and Illumina DNA/RNA UD indexes at the Emory NPRC Genomics Core. Final amplified libraries were purified using AMPure XP bead cleanup and validated by capillary electrophoresis on a TapeStation 4200 (Agilent) before being pooled at equimolar concentrations and sequenced with PE100 reads on an Illumina NovaSeq X Plus and a NovaSeq 6000. A minimum depth of ~2,000,000 reads per sample was targeted.

#### SHAPE-MaP data analysis

Raw FASTQ files were demultiplexed then analyzed using ShapeMapper2 (27) with default parameters and a minimum read depth of 5000. All ShapeMapper2 runs passed the built-in quality control checks. Replicate normalized reactivity values obtained from ShapeMapper2 were averaged and mapped on the 16S rRNA secondary structure (nucleotides C1367– A1542) using RNAcanvas (28). SHAPE reactivities of the partial and non-substrate Δ*rsmH/I/F*, Δ*rsmE/F* Δ*rsmI/E/F*, Δ*rsmH/E/F*, and Δ*rsmH/I/E/F* 30S subunits were compared to Δ*rsmF* 30S subunit (optimal RsmF substrate) using both difference analysis, similarly to as we have done before (29,30), and deltaSHAPE (31). For difference SHAPE analyses, the per-nucleotide averaged normalized SHAPE reactivity values of the Δ*rsmF* 30S were subtracted from those of each other 30S species, with nucleotide reactivity difference classified as strongly decreased (≤ −0.50), moderately decreased (−0.49 to −0.20), no change (−0.19 to 0.19), moderately increased (0.2 to 0.49), and strongly increased (≥ 0.50). For deltaSHAPE, averaged normalized SHAPE reactivity profiles were used as the input for the deltaSHAPE pipeline integrated into the RNAvigate suite (32), with the significance threshold set to the default 1.0. Δ*rsmE/F* − Δ*rsmF* 30S deltaSHAPE analysis was additionally analyzed at a significance threshold of 0.725. Nucleotides with enhanced or protected reactivity indicated by the deltaSHAPE analysis were manually mapped on to the 16S rRNA secondary structures as transparent shaded regions using Adobe Illustrator.

### Molecular dynamics (MD) simulations

#### System set-up

To render MD simulations computationally tractable while preserving the structural context of the ribosomal environment, a partially restrained “ribosomal section” approach was employed as we and others have used previously (33-35). Coordinates corresponding to the ribosomal section system for simulation were extracted from PDB 9Q87 (36) and comprised seven non-contiguous 16S rRNA segments (chain A; nucleotides 6–35, 663–972, 1051–1109, 1189–1236, 1338–1349, 1379–1422, and 1478–1533) and six 30S ribosomal proteins: uS5 (chain E; residues 10–165), uS7 (chain G; residues 2–154), uS11 (chain K; residues 13–129), uS12 (chain L; residues 2–124), uS9 (chain I; residues 111–130), uS10 (chain J; residues 46–66), and bS21 (chain U; residues 2–70). In this starting configuration, 16S rRNA nucleotides A1492 and A1493 occupy a “flipped-in” conformation within the A-site. The included 16S rRNA carries nine intrinsic post-transcriptionally modified nucleotides present in the native *E. coli* 30S subunit arising from the four modifications incorporated by RsmE, RsmF, RsmH and RsmI, as well as those placed by RsmD (m^2^G966), RsmB (m^5^C967), RsmC (m^2^G1207), RsmG (m^7^G1516), and RsmA (m_2_^6^A1518 and m_2_^6^A1519). The fully modified rRNA constituted the wild-type ribosome segment system, with six methylation-depleted variant systems generated by computationally removing the relevant chemical groups. For consistency with descriptions elsewhere, these systems are named for the enzymes that incorporate the removed methyl groups: *ΔrsmF, ΔrsmE/F, ΔrsmI/E/F, ΔrsmH/E/F, ΔrsmH/I/F* and *ΔrsmH/I/E/F*. The remaining six modified nucleotides were retained in all seven systems.

#### MD simulation protocol and analysis

Structure preparation was performed within the Schrödinger Maestro environment, including the addition of hydrogen atoms, bond order assignment, restrained energy minimization, and hydrogen bond optimization. Each system was solvated in a TIP3P water box with 150 mM NaCl. rRNA nucleotides in and near h44 modification sites (9–25, 779–803, 885–934, 1381–1419, and 1481–1530) were unrestrained permitting unbiased sampling of the conformational response to changes in methylation state. All other included 16S rRNA nucleotides and all ribosomal proteins were assigned a light positional restraint (*k* = 0.5 kcal mol^-1^Å^-2^) to maintain overall structural integrity without over-constraining the system. All MD simulations were carried out using the Desmond module of the Schrödinger software suite (v2026-1) with the OPLS4 force field (33,34,37,38).

Each system was subjected to the standard Desmond relaxation protocol prior to production: steepest-descent minimization (up to 2000 steps, convergence criterion 25 kcal mol^-1^Å^-1^), followed sequentially by a 12 psNVT simulation at 10 K with heavy-atom restraints (*k* = 50 kcal mol mol^-1^Å^-2^; Berendsen thermostat), a 12 ps NPT simulation at 10 K and 1 atm with the same restraints (Berendsen thermostat and barostat), a 24 ps NPT simulation at 300 K and 1 atm with heavy-atom restraints, a 24 ps NPT simulation at 300 K and 1 atm with backbone-only restraints, and a 240 ps unrestrained NPT simulation at 300 K and 1 atm (Nosé–Hoover thermostat; Martyna–Tobias–Klein barostat). Each system was subsequently equilibrated for 10 ns in the NPT ensemble (310.15 K, 1 atm). Production simulations of 100 ns for each system were then performed in triplicate in the NPT ensemble (310.5 K and 1 atm; Nosé– Hoover thermostat with relaxation time of 1.0 ps, Martyna–Tobias–Klein barostat with relaxation time of 2.0 ps, and isotropic pressure coupling). Equations of motion were integrated with a multiple time-step scheme: 2 fs for short-range interactions and 6 fs for long-range interactions, with 9 Å nonbonded cutoff. Coordinates were saved every 50 ps.

Simulation quality was assessed using the Desmond simulation quality analysis module. For post-simulation analysis, a total of 23 inter-atomic distances were tracked to assess internal h44 integrity (Watson-Crick base pairing) and cross-strand/ A-site geometry (d2-d6, d8, d10, and d12-d15), h44 inter-helix tertiary contacts with h24 (d19), h31 (d1) and h45 (d7, d9, d11, and d16-d19), and h45 internal cohesion (d21-d23). We also derived 13 pseudo-dihedral angles, defined by N1/N9(n) - C3′(n) - C4′(n+1) - N1/N9(n+1), at m^4^Cm1402 (pseudo-dihedrals *a* and *b*) and m^5^C1407 (pseudo-dihedrals *c* and *d*) on the target strand, and from G1491 to A1500 on the complementary strand (pseudo-dihedrals *e*-*m*). All distances and angles were averaged across the three replicates for each system, with mean and standard deviation values plotted as heat maps using GraphPad Prism 11.

## RESULTS

### Generation of *E. coli* 16S rRNA (h44) methyltransferase deletion strains

To enable studies of the collective impact(s) and functional role(s) of the four 16S rRNA h44 methylations incorporated by RsmH, RsmI, RsmE, and RsmF (**Fig. 1A**), we created a collection of *E. coli* deletion strains lacking two, three or all four of the genes encoding these enzymes. Starting from single gene deletion strains available from the Keio collection (*rsmE, rsmF*, and *rsmH*) (19) or generated previously in our lab (*rsmI*) (20), we first used P1 phage transduction (21) to create the double deletion *E. coli* Δ*rsmH/I*, similarly to what was done before (5), and Δ*rsmE/F* strains (**Fig. 1B**). This process was then repeated to make four triple *E. coli* deletion strains and finally culminating in the generation of the Δ*rsmH/I/E/F* quadruple deletion strain (**Fig. 1B**). The strains all originate from the *E. coli* BW25113 background (19) (used for preparation of “wild-type” 30S subunits), and their deletion genotypes were verified using PCR (**Supplementary Fig. S1A**) and whole genome sequencing (deposited at NCBI BioProject database, accession code PRJNA1482076).

Studies to more broadly determine the phenotypic impacts of the multiple Rsm gene deletions are currently on-going. Interestingly, however, loss of multiple or all four h44 methyl groups does not appear to significantly impact growth compared to wild-type BW25113 at 37 °C, as assessed using a spot dilution assay (**Supplementary Fig. S1B**). In the present study, the availability of 30S subunits hypomethylated at all four sites from the quadruple deletion strain first allowed us to address the question of whether the activity of any of the h44 methyltransferases depends on prior modification(s) at another site.

### RsmF cannot methylate a 30S subunit with fully hypomethylated h44 *in vitro*

Using purified recombinant RsmH, RsmI, RsmE and RsmF, we first confirmed using an *in vitro* [^3^H]-SAM methyltransferase assay that each enzyme can methylate 30S subunits isolated from strains lacking their respective modifications, but not 30S from wild-type *E. coli* BW25113 with a full complement of h44 modifications (**Supplementary Fig. S1C**). Next, using the same assay, but assessed over a 90-minute time course, we compared the activity of each enzyme on 30S subunits lacking only their corresponding modification versus the fully hypomethylated 30S subunit isolated from the quadruple Δ*rsmH/I/E/F* strain (**Fig. 2**).

**Fig. 2.**
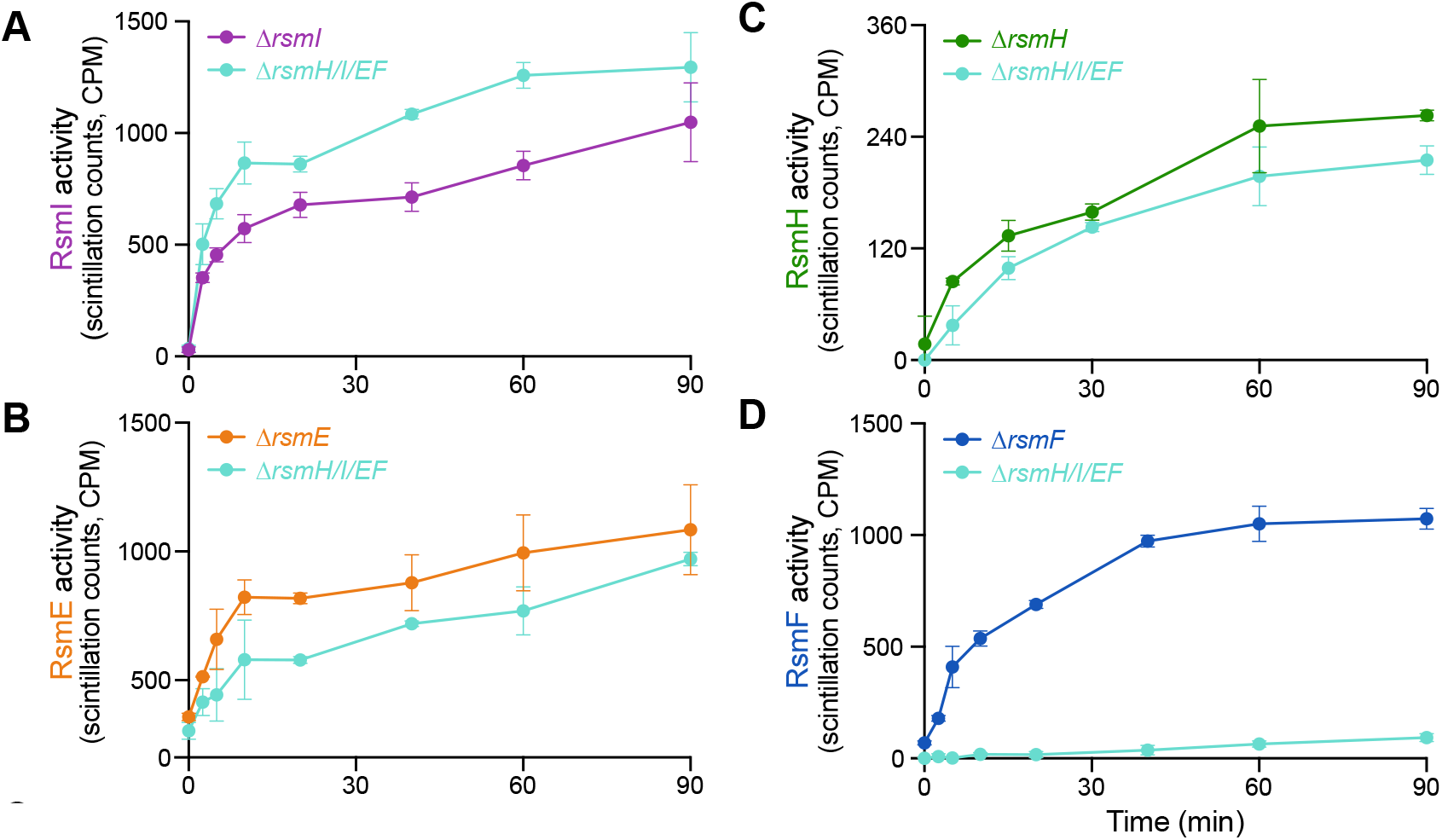
RsmF does not methylate 30S subunits lacking all four h44 methylations. *In vitro* methyltransferase activity assays comparing enzyme activity on 30S subunits isolated from single deletion strains and the quadruple (Δ*rsmH/I/E/F)* strain for (**A**) RsmI (purple), (**B**) RsmE (orange), (**C**) RsmH (green), and (**D**) RsmF (blue). Data for modification of 30S subunits from single deletion strains are color-coded to the corresponding enzyme and those for the Δ*rsmH/I/E/F* are shown in cyan in each plot.

RsmE (m^*3*^U1498), RsmH (m^4^C1402), and RsmI (Cm1402), all modify the partially and fully hypomethylated 30S subunits similarly, but with minor differences: RsmI exhibits slightly higher activity in the absence of the other h44 methyl groups, while RsmE appears to modify the fully hypomethylated 30S subunit slightly less efficiently (**Fig. 2A, B**). Interestingly, RsmH displays essentially the same activity on both substrates, but has much lower activity compared to the other enzymes (**Fig. 2C**). Considering that 30S subunits from wild-type *E. coli* appear to be fully methylated at m^4^C1402 (**Supplementary Fig. S1C**), it is possible that the set-up or time-frame of our *in vitro* assay does not fully reflect substrate accessibility to RsmH in the cell. Alternatively, a cellular component needed by RsmH may be absent *in vitro*, but this possibility will require further investigation. In sharp contrast to the other three enzymes, RsmF (m^5^C1407) shows essentially no activity over the duration of the time-course in the absence of the other h44 methyl groups (**Fig. 2D**), indicating that RsmF depends on the prior action of one or more of the other h44 methyltransferases.

### RsmF activity requires prior h44 modification in the *E. coli* cell

To confirm that the unanticipated inability of RsmF to *in vitro* methylate the Δ*rsmH/I/E/F* 30S is not an artifact of the sample preparation process or the assay itself, we next asked whether RsmF activity is similarly impeded by the absence of other h44 modifications in *E. coli* using 16S rRNA bisulfite sequencing. This approach uses sodium bisulfite to deaminate all non-m^5^C cytosine nucleotides (to uracil), resulting in T and C sequencing reads for unmodified and modified positions, respectively.

For comparisons of m^5^C1407 modification in distinct h44 hypomethylation contexts, 16S rRNA was extracted from 30S subunits isolated from wild-type BW25113 and the Δ*rsmF*, Δ*rsmH/I/E* and Δ*rsmH/I/E/F* strains and treated with bisulfite reagent. The 30S subunits from the wild-type and quadruple deletion strains serve as controls for fully modified and fully hypomethylated h44, respectively, while comparison of Δ*rsmF* 30S and Δ*rsmH/I/E* informs on the requirement of modifications by RsmE, RsmH and/or RsmI on the activity of RsmF in the cell. Treated 16S RNA was first converted to a ~100 nucleotide double-stranded DNA amplicon encompassing C1407 (and C1402) by reverse transcription and cDNA PCR. As an independent quality control check on the modification process, we similarly amplified the region surrounding C967 (modified to m^5^C by RsmB), which should be unchanged in 30S subunits from all strains. To account for potential variability in extent of 16S rRNA modification and bisulfite reaction efficiency, dsDNA amplicons were cloned and sets of 5 (C967) or 10 (C1407) individually isolated plasmids corresponding to unique converted 16S rRNAs were sequenced. Using the resulting unambiguous trace files, the full extent of the *in vivo* methylation pattern differences among the strains could be assessed. As expected, full protection of C967, but not the surrounding cytosine nucleotides, was observed in each case (**Supplementary Fig. S2**), indicating successful bisulfite conversion and that this distant site is fully modified in all strains with no unanticipated effects of the h44 methyltransferase gene deletions.

Within the region encompassing C1407 (N = 10), for wild-type 16S rRNA we observe complete and high protection at C1407 (0% converted) and C1402 (20% converted), respectively, matching previous analyses (24) (**Fig. 3A**). C1402 protection arises due to m^4^C base modification by RsmH but does not appear as efficiently protected as for m^5^C, likely due to differences in methylation placement. As anticipated, Δ*rsmF* 16S rRNA retains near complete protection at C1402 (30% converted), but with almost no protection at C1407 (90% converted), consistent with both the lack of RsmF activity and the ability of the remaining three enzymes to modify the 30S subunit in the absence of RsmF. (**Fig. 3B**). Similarly, Δ*rsmH/I/E/F* 16S rRNA shows the expected complete lack of protection at C1402 and C1407 (both 100% conversion), consistent with the lack of RsmH and RsmF activity in this strain (**Fig. 3C**). Finally, the Δ*rsmH/I/E* 16S rRNA shows the expected lack of protection at C1402 (100% conversion) due to the absence of RsmH, but also displays substantially reduced protection at C1407 (70% conversion) despite encoding RsmF (**Fig. 3D**). To verify this result, a second independent set of 10 colonies was analyzed, resulting in the same 70% conversation rate at C1407. These findings reveal that the ability of RsmF to modify the 30S subunit within the cell is significantly hindered in the absence of the other three h44 methyltransferases, consistent with the *in-vitro* methylation results (**Fig. 2D**).

**Fig. 3.**
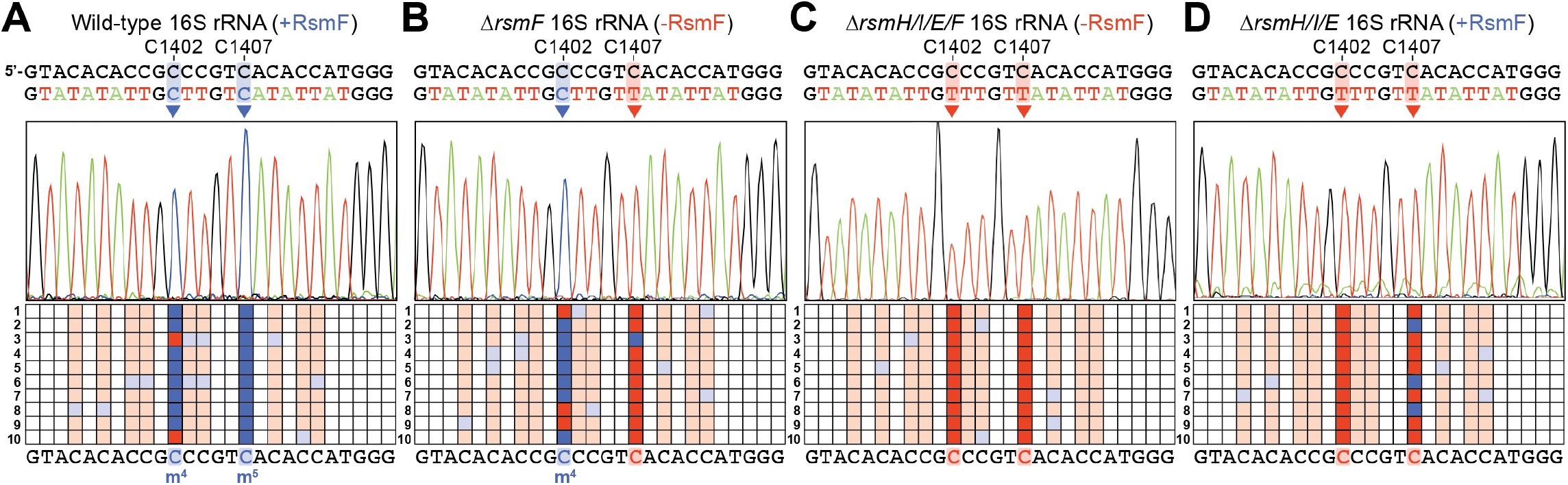
C1407 methylation by RsmF in cells is reduced in the absence of the other h44 methyltransferases. RNA bisulfite sequencing analysis of the 16S rRNA h44 region surrounding C1402 and C1407 for 30S subunits isolated from (**A**) wild-type BW25113, (**B**) Δ*rsmF*, (**C**) Δ*rsmH/I/E/F*, and (**D**) Δ*rsmH/I/E*. In each panel, the 16S rRNA sequence is shown (*top*) with the sequencing color-coded bisulfite converted sequence below. Also shown are representative sequencing chromatograms (*middle*) and grid maps summarizing C to T conversion at each cytosine position in the individual sequenced clones 1-10 (*bottom*). Converted sites (C to T) are shown in red and unconverted sites in blue, with the C1402 and C1407 positions indicated and highlighted with the dark shades of each color.

### RsmF activity specifically requires the modifications incorporated by RsmE and RsmH

Having established the dependency of RsmF on prior deposition of one or more of the other h44 modifications, we next set out to determine which specific h44 methylation(s), or corresponding enzyme(s), are required for RsmF activity. To this end, we returned to the *in vitro* assay to compare the activity of RsmF on 30S subunits lacking C1407 methylation and various combinations of the other h44 methylations. (**Fig. 4A**). Absence of U1498 methylation (Δ*rsmE/F* 30S) or both C1402 modifications (Δ*rsmH/I/F* 30S) each reduced modification of C1407 by RsmF, with the former having the largest effect, with 37% and 63% activity compared to Δ*rsmF* 30S, respectively. Absence of both RsmI (Cm1402) and RsmE (m^3^U1498) inΔ*rsmI/E/F* 30S further decreases RsmF activity to 16% compared to Δ*rsmF* 30S. However, only the combined absence of RsmH (m^4^C1402) and RsmE (m^3^U1498) fully recapitulates the complete ablation of RsmF activity already observed with the fully hypomethylated substrate (compare *rsmH/E/F* 30S and Δ*rsmH/I/E/F* 30S with *rsmF* 30S). Thus, even though each of the h44 methyltransferases appears to influence RsmF activity, it is the combined presence of both RsmH and RsmE that is necessary for RsmF to modify its substrate.

**Fig. 4.**
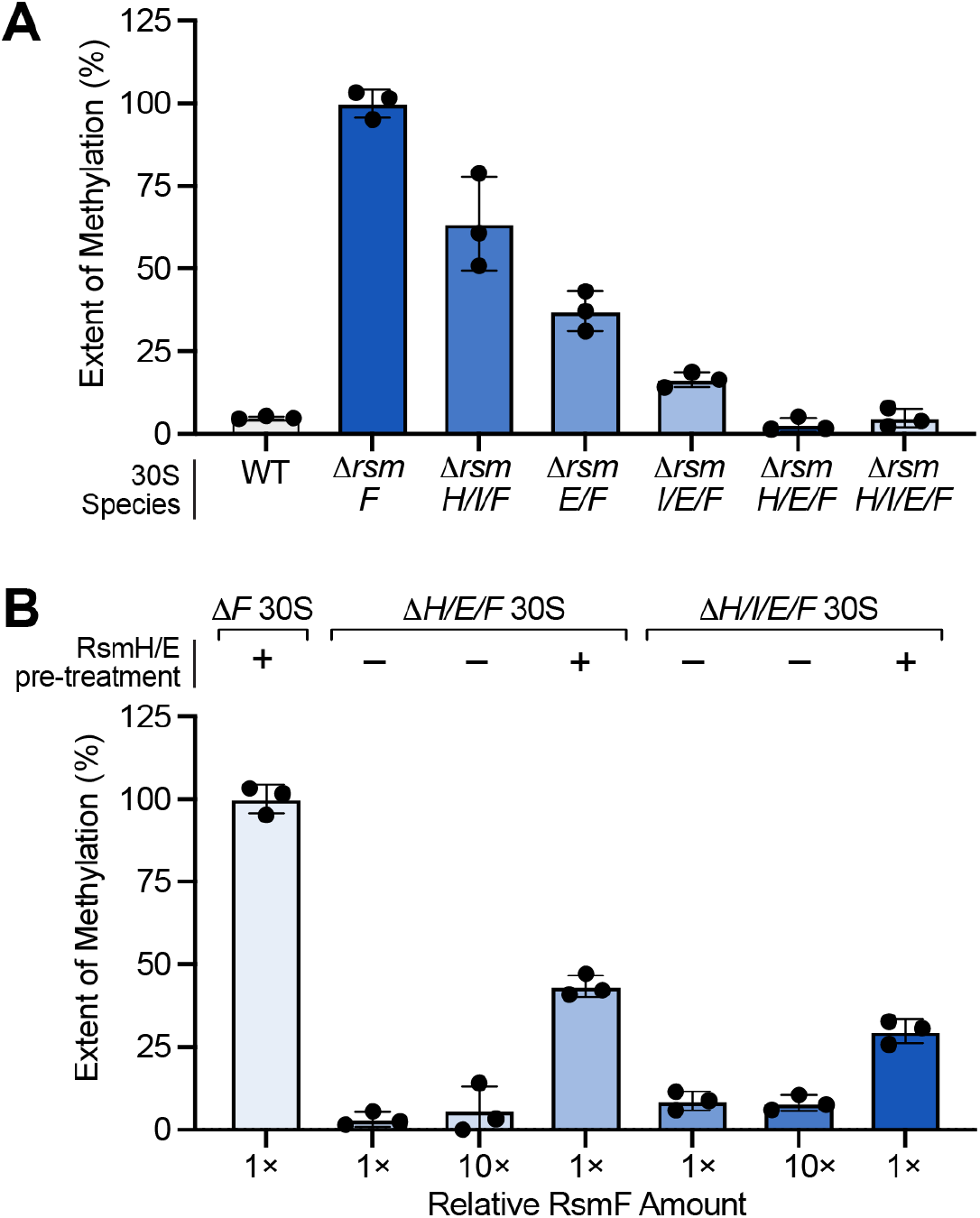
RsmF depends on prior deposition of the methyl groups by RsmE and RsmH. *In vitro* methyltransferase activity assays evaluating RsmF enzyme activity on 30S ribosomal subunits. (**A**) Relative extent of RsmF enzyme activity on 30S subunits isolated from Δ*rsmF*, Δ*rsmH/I/F*, Δ*rsmE/F*, Δ*rsmI/E/F*, Δ*rsmH/E/F*, Δ*rsmH/I/E/F*, and wild-type (WT) strains. Activity is expressed as a percentage relative to Δ*rsmF* substrate. (**B**) RsmF activity on unmodified versus sequentially modified 30S subunits. Substrates isolated from Δ*rsmF*, Δ*rsmH/E/F*, and Δ*rsmH/I/E/F* strains were evaluated either with (+) or without (−) prior enzymatic pre-treatment by RsmH and RsmE. Assays were conducted using either standard (1×) or a ten-fold excess (10×) relative concentration of RsmF. Activity is expressed as a percentage relative to the treated Δ*rsmF* substrate. Bar graphs represent the mean percentage of methylation.

Whether the dependency of RsmF upon RsmE/RsmH arises directly from the deposited methyl groups and their influence on 16S rRNA structure or stability, or upon some other facet of these enzymes’ prior action on the 30S subunit cannot be discerned from the results thus far. For example, as 16S rRNA methyltransferases functioning late in 30S assembly, RsmH and RsmE likely need to manipulate a substantially folded 30S structure to access their target nucleotides, as previously observed for RsmI (20). Thus, the observed dependency of RsmF could arise from the requirement for prior binding and specific unfolding of the 16S rRNA by RsmH and RsmE. To test this possibility, we designed catalytically inactive variants of RsmH (D55R) and RsmE (G194R) that cannot bind the co-substrate SAM, but which retain all residues necessary to interact with the 30S subunit. After confirming that these variants are catalytically inactive and do not interfere with RsmF activity on its favored substrate (Δ*rsmF* 30S), we mixed each enzyme with RsmF individually and combined to assess whether their presence could restore its activity on Δ*rsmH/E/F* 30S (**Supplementary Fig. S3**). However, no combination of either RsmH (D55R) or RsmE (G194R) was found to rescue RsmF activity.

To test the second possibility–that it is the presence of the methyl groups that is required for RsmF activity–we pre-treated Δ*rsmH/E/F* 30S, Δ*rsmH/I/E/F* 30S, and, as control, Δ*rsmF* 30S with wild-type RsmH and RsmE using non-radiolabeled SAM. After removal of the enzymes and SAM, the *in vitro* modified 30S subunits were then used in subsequent assays with [^3^H]-SAM. As anticipated, the activity of both RsmH and RsmE is drastically decreased on these substrates pre-treated at their target sites compared to their non-treated counterparts, even with excess (10×) enzyme, confirming essentially complete pre-methylation (**Supplementary Fig. S4A-D**). Additionally, both RsmI and RsmF display similar activity on pre-treated Δ*rsmH/I/E/F* 30S and Δ*rsmF* 30S, respectively, compared to “mock”-treated versions that underwent the treatment but without prior addition of RsmH and RsmE (**Supplementary Fig. S4E-F**). However, both RsmI and RsmF exhibit some decrease in activity (~30%) with their respective mock and pre-treated substrates compared to the non-treated versions (**Supplementary Fig. S4E-F**) indicating that the steps to remove pre-treatment enzymes and SAM cause some “damage” to the 30S subunits generally reducing their suitability as modification substrates. Nonetheless, these results show that the pre-treated 30S subunits are suitable substrates to test for restoration of RsmF activity with placement of the m^4^C1402 (RsmH) and m^3^U1498 (RsmE) modifications. Indeed, we find that RsmF activity is restored to ~ 43% and ~ 30% that of the pre-treated Δ*rsmF* 30S control for Δ*rsmH/E/F* and Δ*rsmH/I/E/F* 30S, respectively. In contrast, RsmF could not methylate either of the two 30S subunits without RsmH/RsmE pre-treatment, even at 10 times higher relative enzyme concentration (**Fig. 4B**). Collectively, these studies thus reveal that it is the presence of the m^4^C1402 and m^3^U1498 modifications incorporated by RsmH and RsmE, respectively, that are required for RsmF activity.

### Absence of h44 methylations required by RsmF alters nucleotide dynamics opposite C1407

To gain deeper insight into how the h44 methyl groups deposited by RsmH and RsmE might control the ability of RsmF to modify its target site, we turned to selective 2′-hydroxyl acylation analyzed by primer extension and mutational profiling (SHAPE-MaP) to assess the local nucleotide environment. Using the series of hypomethylated 30S subunits assessed earlier and which support a range of RsmF activities from viable to non-substrates (Δ*rsmH/I/F* 30S > Δ*rsmE/F* > Δ*rsmI/E/F* > Δ*rsmH/E/F* ~ Δ*rsmH/I/E/F* 30S; **Fig. 4A**), we specifically focused on *in vitro* SHAPE reactivities of h44 and h45 (nucleotides 1367-1542; **Supplementary Fig. S5**) and compared how these reactivities differ from those of the preferred RsmF substrate, Δ*rsmF* 30S which lacks only the m^5^C1407 methylation. Two approaches were used to compare reactivities with the Δ*rsmF* 30S: Difference SHAPE, in which the averaged normalized SHAPE reactivities for each nucleotide were subtracted, and deltaSHAPE (31), which accounts for the associated errors in the reactivity differences between species as well as the relative magnitude of differences to provide a more statistically rigorous approach (**Supplementary Fig. S6**). The results of both analyses were mapped onto the 16S rRNA secondary structure of h44 and h45 (**Fig. 5**).

**Fig. 5.**
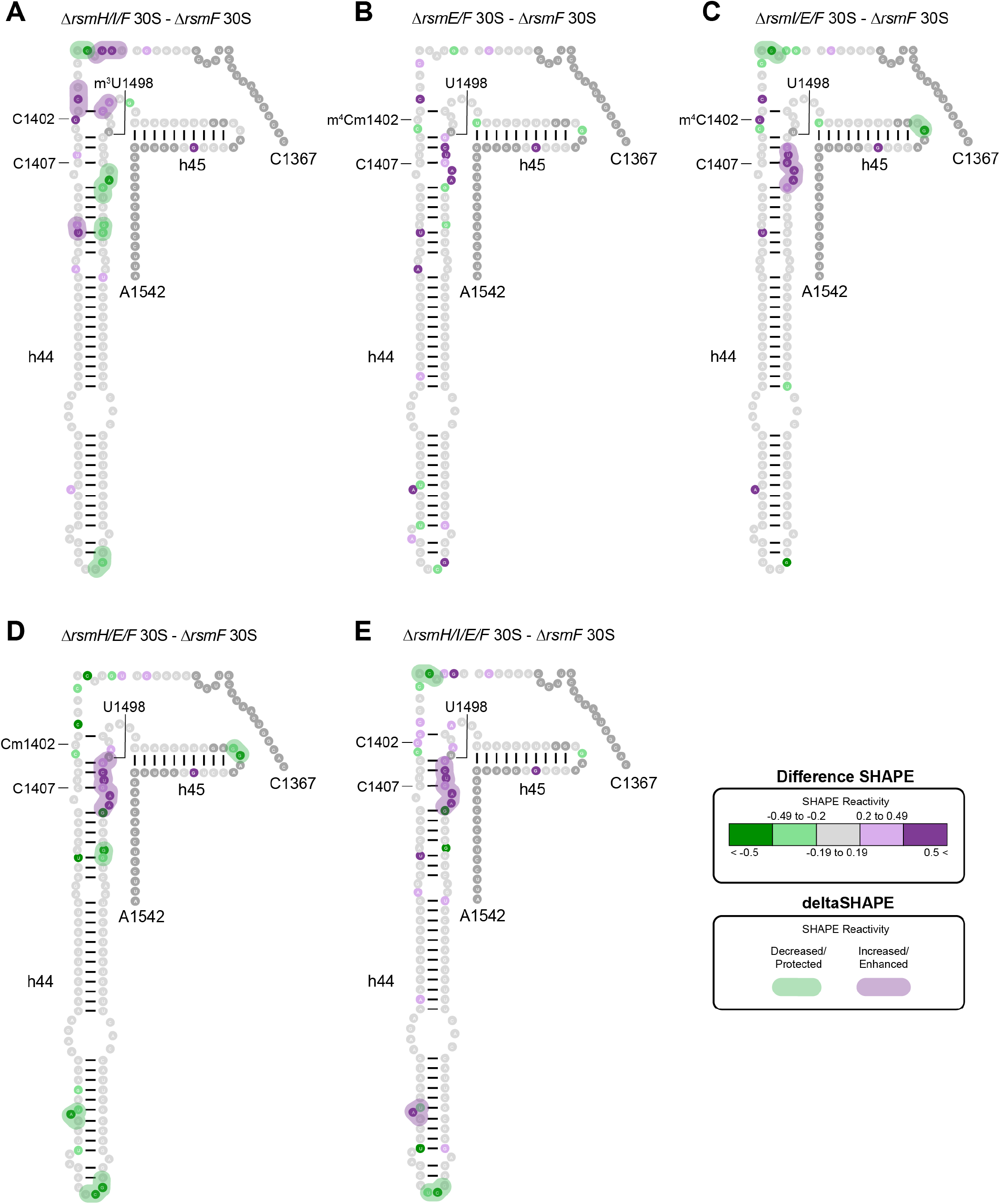
h44 nucleotide dynamics opposite the RsmF target site likely dictate loss of activity on hypomethylated 30S. *In vitro* SHAPE reactivities of the 16S rRNA region encompassing h44 and h45 (nucleotides 1367–1542). Each secondary structure shows the SHAPE-MaP reactivity profiles after subtraction of values for Δ*rsmF* 30S from (**A**) Δ*rsmH/I/F* 30S, (**B**) Δ*rsmE/F* 30S, (**C**) Δ*rsmI/E/F* 30S, (**D**) Δ*rsmH/E/F* 30S, and (**E**) Δ*rsmH/I/E/F* 30S. Differences are represented using both Difference SHAPE (individually colored circles) and deltaSHAPE (background shading) with green and purple indicating decreased and increased differential SHAPE reactivities, respectively.

Compared to the Δ*rsmF* 30S, loss of dual C1402 methylation by RsmH and RsmI in the Δ*rsmH/I/F* 30S causes alterations in the SHAPE reactivity of the nucleotides bridging h44 to h43 and to h45, as well as a decrease in the reactivity of h44 nucleotides U1491–A1493 in the strand across from C1407, and these differences are roughly corroborated by both Difference SHAPE and deltaSHAPE (**Fig. 5A** and **Supplementary Fig. S6A**). However, since, the Δ*rsmH/I/F* 30S is still a viable substrate for RsmF (**Fig. 4A**), it is not possible to specifically discern whether such changes in nucleotide dynamics are critical for RsmF. In contrast, loss of m^3^U1498 methylation by RsmE in the Δ*rsmE/F* 30S predominantly results in increased SHAPE reactivity of h44 nucleotides A1492–G1497 as assessed by Difference SHAPE calculations (**Fig. 5B**). This increase in h44 nucleotide flexibility opposite C1407 may explain why RsmF activity is much more perturbed with the Δ*rsmE/F* 30S compared to the to the Δ*rsmH/I/F* 30S (**Fig. 4A**). Notably, while these changes were not identified via deltaSHAPE at the initial significance threshold (1.0), corresponding changes can be observed with a lower significance threshold of 0.725 (**Supplementary Fig. S7**). In the absence of one of the C1402 methylations by RsmI or RsmH in addition to m^3^U1498 methylation, in the poorer RsmF substrates Δ*rsmI/E/F* and Δ*rsmH/E/F* 30S, respectively, this increased reactivity opposite C1407 becomes more pronounced and is identified by both approaches, including at the higher threshold for deltaSHAPE (**Fig. 5C-D** and **Supplementary Fig. S6B-C**).

Comparing how the Δ*rsmI/E/F* 30S (significantly diminished RsmF activity) and Δ*rsmH/E/F* 30S (no RsmF activity) differ in their SHAPE-MaP profiles to Δ*rsmF* 30S, we notice similar effects of decreased reactivity of h45 nucleotides G1516-G1517 and increased reactivity of h44 nucleotides G1491-C1496, again supported by both Difference and deltaSHAPE (**Fig. 5C-D**). However, loss of RsmH methylation appears to extend the range of the h44 nucleotides opposite C1407 experiencing increased reactivity (G1491–A1499; +/− 1 nucleotide depending on the method of analysis) not observed with loss of RsmI methylation. This increased number of h44 reactive nucleotides could also be observed with the Δ*rsmH/I/E/F* 30S, which generally showed a difference profile that appeared as a combination of all the others (**Fig. 5E** and **Supplementary Fig. S6D**).

In summary, loss of U1498 methylation by RsmE appears to have a destabilizing effect on the h44 nucleotides adjacent to it, on the h44 strand opposite C1407. Loss of either of the C1402 methyl groups appears to further strengthen the destabilizing effect, with m^4^C1402 methylation by RsmH increasing the range of nucleotides experiencing increased flexibility. The h44 strand opposite C1407 thus appears critical for RsmF to recognize its substrate for modification, with increased flexibility potentially impeding RsmF binding and thereby explaining the necessity of prior methylation by RsmH and RsmE for its activity.

### MD simulations reveal distinct roles for m^3^1498 and m^4^C1402 in shaping rRNA structure and dynamics

One limitation of SHAPE-MaP is that it requires the ribose 2’-OH to be dynamic and solvent accessible for reaction with the SHAPE probe. Additionally, because RsmF requires a mature 30S substrate in vitro, it likely recognizes a three-dimensional 16S rRNA surface to bind to its substrate, but our SHAPE analysis did not provide information about structural changes between helices. Thus, to gain further atomic insight how h44 methylation status shapes the decoding center for RsmF recognition, all-atom MD simulations were performed on a ribosomal section (extracted from PDB 9Q87), encompassing h44, with A1492/A1493 in a flipped-in orientation, and neighboring helices h24, h31 and 45. In total, seven distinct systems were simulated, including fully methylated and six hypomethylated RsmF substrates (ranging from viable to non-substrates for RsmF). Structural dynamics in each system were quantified using 23 inter-atomic distances, assessing Watson-Crick within h44 as well as multiple inter-helix contacts, and 13 pseudo-dihedral angles, defined in the Materials and Methods, spanning both strands of h44 (**Fig. 6A**).

**Fig. 6.**
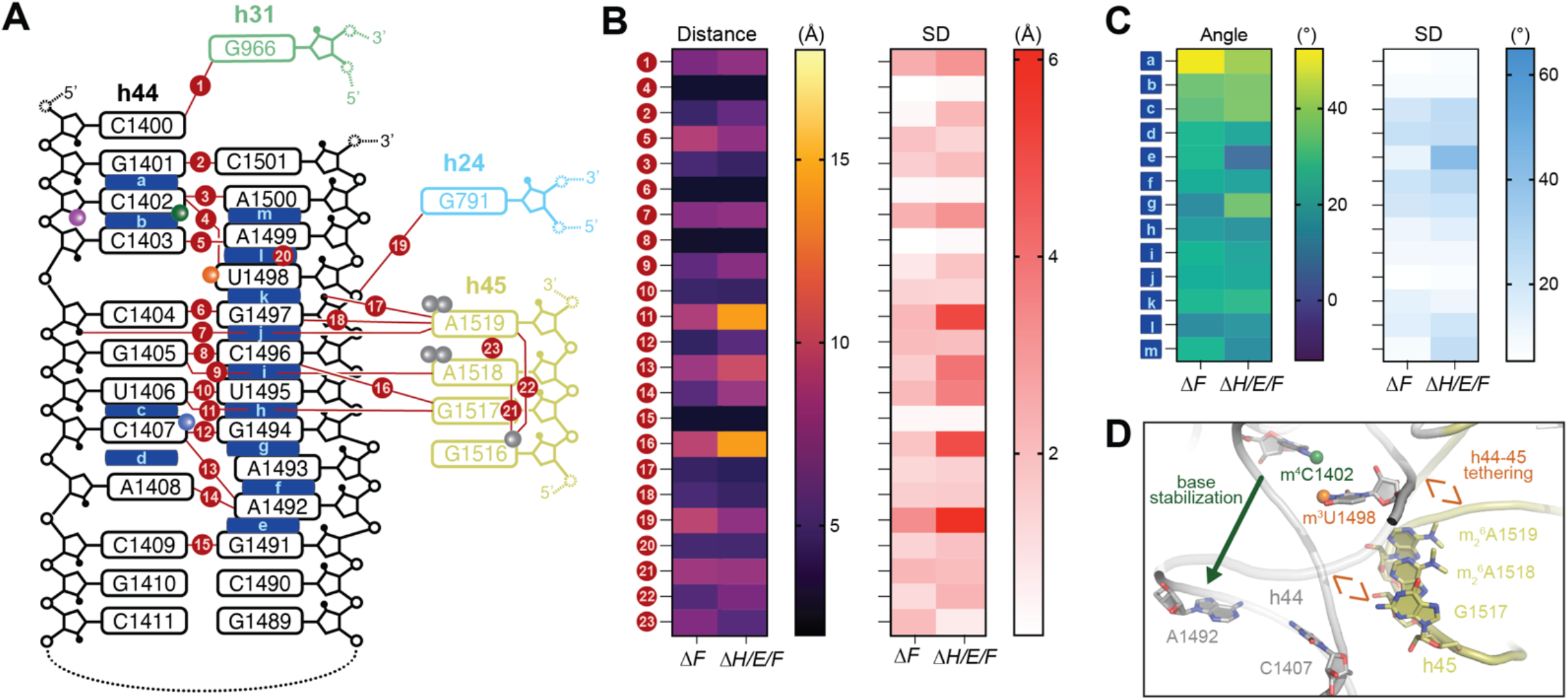
Loss of m^3^1498 and m^4^C1402 methylation destabilizes h44 and its interaction with h45. (**A**) Schematic depicting the inter-atomic distances (numbered) and pseudo-dihedral angles (lettered) measured to assess the impact of hypomethylation on h44 structure and rRNA-rRNA interactions. Comparison of the Δ*rsmF* 30S and Δ*rsmH/E/F* 30S simulated systems shown as: (**B**) Heat map representation of interatomic distances (*left*) and accompanying standard deviations (*right*); and (**C**) heat map representation of the pseudo-dihedral angles (*left*) and accompanying standard deviations (*right*). (**D**) model (derived from PDB 9Q87) summarizing the main insights from the MD analysis: h44-45 tethering by m^3^U1498 and A1492 base stabilization by m^4^C1402.

The simulated Δ*rsmF* 30S largely resembles the fully methylated system in all assessed metrics, indicating that loss of m^5^C1407 methylation alone does not cause substantial perturbation. Comparing substrate Δ*rsmF* 30S and non-substrate Δ*rsmH/E/F* 30S, two major trends are apparent in the h44/h45 interaction and in the base stacking within h44. First, the distances assessing the h44-h45 junction interaction between U1406 and G1517 (d11: Δ*H/E/F* 14.0 ± 5.2 Å and Δ*F* 8.6 ± 2.1 Å) and between C1496 and G1517 (d16: Δ*H/E/F* 13.9 ± 5.0 Å and Δ*F* 9.2 ± 1.8 Å) are significantly higher in Δ*rsmH/E/F* 30S, with increased dynamics as assessed by the increased standard deviation (SD) of both parameters (**Fig. 6B**). These observations indicate that h45 is pulling away from h44 in the hypomethylated system. Similar increases in h44-h45 junction distances could be observed in all other systems lacking m^3^U1498 methylation, but not in Δ*rsmH/I/F* 30S (**Supplementary Fig. S8A**,**B**). Because of the position of m^3^U1498 at the h44/h45 junction, it likely makes a direct van der Waals contribution to h44/h45 packing, loss of which could untether h45 from h44. The destabilization of h44/h45 inter-helix contact may thus lead to the increased flexibility of the 3’ strand of h44 (A1492–G1497 ± 1 nucleotide) that we observe in our SHAPE-MaP analyses (**Fig. 5**), offering further insight as to why RsmF exhibits poor or no activity on 30S subunit substrates lacking m^3^U1498 (**Fig. 4A**).

The second trend from comparison of the Δ*rsmF* and Δ*rsmH/E/F* 30S systems is the collapse of the pseudo-dihedral angle between G1491–A1492 (**Fig. 6A**, *dihedral e*), with a substantial decrease in the mean angle (Δ*H/E/F* 6.2° and Δ*F* 30.6°) accompanied by a dramatic increase in variability (SD Δ*H/E/F* 41.9° compared to Δ*F* 12.6°) (**Fig. 6C**). This change is correlated with additional more modest increases in cross-strand distances between A1492 and C1407 (d13: Δ*HE/F* 10.0 ± 4.0 Å and Δ*F* 7.9 ± 1.5 Å) and between A1492 and A1408 (d14: Δ*H/E/F* 7.8 ± 3.6 Å and Δ*F* 5.3 ± 2.1 Å) (**Fig. B**). The SD of the pseudo-dihedral angle serves as a marker for the heterogeneity of geometries sampled by a base, indicating that A1492 exhibited higher heterogeneity in its orientation in Δ*H/E/F* 30S. We only observe similar increases in SD of pseudo-dihehdral angle between G1491–A1492 (*dihedral e*) in systems missing m^4^C1402 methylation (i.e., Δ*H/I/F* and Δ*H/I/E/F*; **Supplementary Fig. S8C**,**D**), highlighting that RsmH methylation of C1402 helps maintain the rigidity of A1492. In summary, these simulations provide additional, complementary insight into the roles of both the m^3^U1498 and m^4^C1402 as an anchor for h44-45 tethering and stabilizer of A1492, respectively (**Fig. 6D**), thereby priming rRNA surrounding C1407 for modification by RsmF.

## DISCUSSION

Of the 10 methylated nucleotides in the *E. coli* 16S rRNA of the 30S ribosomal subunit, three are clustered in close proximity in h44 around the decoding center. Although there is a growing body of literature highlighting the role(s) of each of the four methyl groups in this region (5,9,12-17,40), the collective effect of the h44 nucleotide methylation cluster on the structure and function of the decoding region has yet to be evaluated. To enable studies addressing this critical gap, we generated a collection of *E. coli* deletion strains lacking various combinations of h44 methyltransferases. Interestingly, our preliminary analyses of these strains revealed that the absence of all four h44 methyl groups does not detectably impact growth under standard laboratory conditions (**Supplementary Fig. S1C**) (41). However, based on earlier studies in yeast of combinatorial large subunit rRNA hypomethylation (42,43) as well as the effects of loss of individual bacterial h44 methylations, we anticipate that that the greatest phenotypic deficits in our new strains will manifest under conditions of stress. Studies investigating these deletion strain phenotypes are currently on-going.

In the current work, we used 30S subunits derived from the deletion strain collection to reveal an unanticipated dependency of the m^5^C1407 methyltransferase RsmF on prior modification of h44 by the other 16S rRNA methyltransferases that act adjacent to the decoding center. *In vitro*, RsmF exhibited a complete inability to modify a mature but fully h44-hypomethylated 30S subunit (**Fig. 2D**). This finding was recapitulated in the *E. coli* cell, where bisulfite sequencing analyses revealed a sharp reduction (though not complete elimination) of C1407 protection (due to m^5^C modification) in the absence of the other h44 methyltransferases (**Fig. 3D**). One explanation for the discrepancy between the *in vitro* and *in vivo* results could be the nature of the substrates used in both experiments. For the *in vitro* methylation assays, mature 30S subunits are purified from the cell (in absence of RsmF) and incubated with RsmF for the first time for only the limited duration of the assay. In contrast, in the cell, endogenous RsmF (when present) could potentially access its 30S substrate for a longer period during the entirety of the 30S biogenesis process, before rRNA extraction and treatment with bisulfite reagent. Nonetheless, despite potential differences in the timing and duration of enzyme action, as well as in the nature of the substrate, both *in vitro* and *in vivo* approaches highlight a strong dependency of RsmF on prior methylation at adjacent locations in h44.

Through functional assays with 30S subunits containing different combinations of h44 hypomethylation, we specifically demonstrated that the prior action of RsmE (m^3^U1498) and RsmH (m^4^C1402) are necessary for RsmF to deposit its modification on C1407 (**Fig. 4A**). Furthermore, we could de-couple the potential 16S rRNA structural manipulation capacity of these enzymes from the effects the methyl groups themselves directly impart on h44 structure and dynamics (**Fig. 4B** and **Supplementary Fig. S3**). These analyses clearly demonstrate that it is the deposited methyl groups themselves that are necessary for the formation of a 30S subunit substrate on which RsmF can act. While the modifications placed by RsmE and RsmH appear to be the major drivers of this effect, we note that greater recovery of RsmF activity was possible with the pre-RsmE/RsmH treated Δ*rsmH/E/F* 30S than with the Δ*rsmH/I/E/F* 30S (**Fig. 4B**). This suggests that the absence of the Cm1402 modification in Δ*rsmH/I/E/F* 30S, also helps maintain a favorable 16S rRNA structure for RsmF activity, but its presence cannot compensate for the absence of the other modifications at m^4^C1402 and m^3^U1498.

RNA methylation is known to alter the local environment by modifying electrostatic sites, hydration patterns, and enhancing base-stacking interactions (44,45). To probe how the h44 methylations optimally shape the 16S rRNA decoding center region for RsmF to modify its substrate, we assessed their effects on h44 structure and nucleotide dynamics using SHAPE-MaP. This analysis showed that the combined loss of the RsmE-deposited methylation and that of RsmH or RsmI increases the flexibility of the h44 strand directly opposite the C1407 target site. Further, the combined loss of the RsmE/ RsmH modifications has a more pronounced effect, increasing the flexibility of a greater number of h44 nucleotides (**Fig. 5**), consistent with the complete loss of RsmF activity on this hypomethylated 30S subunit but retention of some activity with the Δ*rsmI/E/F* 30S. MD simulations further revealed that loss of the m^3^U1498 modification disrupts the h44–h45 junction, offering a rationale for the increased flexibility in the h44 3’-strand observed by SHAPE. The simulations also showed that the specific loss of C1402 methylation by RsmH, but not by RsmI, contributes to the flexibility of A1492 and its adoption of multiple conformations. Taken together, findings from these complementary approaches offer a compelling model for how RsmF depends on both the m^3^U1498 and m^4^C1402 methyl groups. RsmF likely requires a high degree of rigidity in the complementary h44 strand to its target nucleotide, with A1492 playing a key role in substrate recognition. This rigidity is primarily provided by m^3^U1498 tethering the h44-45 junction and m^4^C1402 stabilizing A1492 (**Fig. 6D**) for RsmF to engage with its target. Nonetheless, high-resolution structural studies will be necessary to fully define the molecular mechanism of substrate recognition and modification by RsmF.

In direct contrast to the dependency of RsmF on intrinsic h44 methylation, previous studies showed that it is inhibited by prior modification of h44 by the acquired aminoglycoside-resistance m^1^1408 methyltransferase NpmA (46). RsmF thus appears particularly sensitive to alterations in methylation patterns around its target site. Based on this observation, and given the directly adjacent locations of their target nucleotides, NpmA and RsmF were proposed to physically clash with each other, with the former able to impede the latter (46). However, our rRNA structure probing studies and insight into the role of methylation-mediated changes in rRNA structure and nucleotide dynamics, may provide an alternative rationale to the inhibition: the m^1^A1408 methyl group itself might similarly induce changes in h44, mirroring the impact of loss of m^3^U1498 and m^4^C1402 methylations, to interfere with RsmF function. Further studies exploring the interplay of intrinsic and acquired aminoglycoside-resistance h44 methylations will be necessary to test this idea.

The dependency of an intrinsic rRNA modifying enzyme on the actions of another has been observed in other regions of the ribosome. For instance, the 16S rRNA m^2^G966 methyltransferase RsmD cannot modify its target until the binding of ribosomal proteins S7 and S19, whose sequential addition is coordinated by the actions of 16S rRNA m^5^C967 methyltransferase RsmB (47,48). Likewise, eukaryotic 18S rRNA N1-methyltransferase Emg1 requires prior psuedouridylation of its substrate nucleotide U1191 by the H/ACA box snoRNP snR35 for its function (49). However, whereas these examples represent an indirect, ribosomal-assembly mediated dependency (in the case of RsmD) or prior modification on the same target nucleotide (in the case of Emg1), our findings reveal RsmF to be currently unique in its reliance on the direct chemical effects of methyl groups at distant sites for its activity.

Overall, our findings offer insight into the timing of modification incorporation during 30S assembly and provide a critical framework for assessing the collective impact of h44 methylation on 30S function. Specifically, we establish that RsmF acts sequentially after RsmH and RsmE, and potentially RsmI, deposit their modifications on h44 late in 30S biogenesis. Why this order of modifications has been maintained in 30S biogenesis is currently unclear. However, the m^5^C1407 modification has been shown to act as a kinetic regulator of the A site, slowing the transition rate between the ground and excited states, and adjusting the timeframe of proper codon-anticodon recognition (16). Additional effects on translation upon loss of m^5^C1407 methylation have also been noted, such as decreases in stop-codon readthrough efficiency (46) and modest increases in ectopic protein expression (15). These observations appear to position methylation of C1407 by RsmF as one of the final steps of 30S subunit assembly, ensuring the proper fine-tuning of the decoding center. This late step may be gated by prior modification of h44 to ensure that it only occurs after the more foundational structural and chemical assembly checkpoints have been reached during 30S biogenesis.

## DATA AVAILABILITY

All strains and corresponding genomic data generated in this study have been deposited in the NCBI BioProject database under accession number PRJNA1482076. Sequencing data for SHAPE-MaP analyses are available through the NCBI Sequence Read Archive (https://www.ncbi.nlm.nih.gov/sra) under BioProject accession number PRJNA1482075. All other data are available in the main text or the supplementary materials.

## SUPPLEMENTARY DATA

Supplementary Data are available at NAR online.

### ACKNOWLEDGMENTS

We thank Dr. Marcin Grabowicz for input and guidance on the process of *E. coli* deletion strain generation.

