## Supplementary Information for "*E. coli* RsmF activity depends on prior modification of 16S rRNA helix 44"

**This PDF file includes:**
- Supplementary Figures S1-S8
- Supplementary Tables S1-S2

### Supplementary Figures

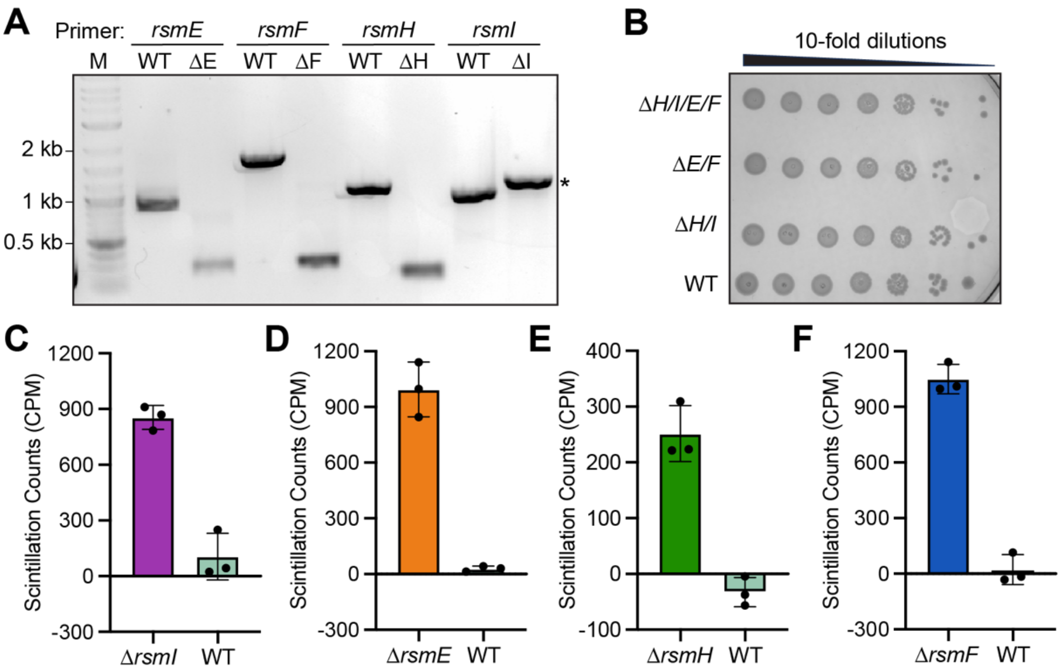


**Fig. S1. Rsm deletion strain verification, viability, and methylation assays.** (**A**) PCR amplification confirming the generation of single-gene knockout strains. Genomic DNA from wild-type (WT) and the deletions Δ*rsmE*, Δ*rsmF*, Δ*rsmH*, Δ*rsmI* was amplified using gene-specific primers. The lower molecular weight bands in the Δ*rsmE*, Δ*rsmF* and Δ*rsmH* mutant lanes confirm the deletion of the target alleles and removal of the selection marker. The higher molecular weight band in the Δ*rsmI* lane (*) indicates the replacement of the native gene with a selectable marker cassette that is larger than the wild-type allele. (**B**) Cell viability and growth assessment via spot dilution assay. Cultures of WT, double-deletions (Δ*rsmH/I* and Δ*rsmE/F*), and the quadruple-deletion strains (Δ*rsmH/I/E/F*) were serially diluted (10-fold) and spotted onto agar to evaluate baseline growth phenotypes. (**C**–**F**) *In vitro* methyltransferase activity assays comparing enzyme activity on their respective 30S subunits (unmethylated only at the single target site) isolated from (**C**) Δ*rsmI*, (**D**) Δ*rsmE*, (**E**) Δ*rsmH*, and (**F**) Δ*rsmF* to fully modified WT controls.

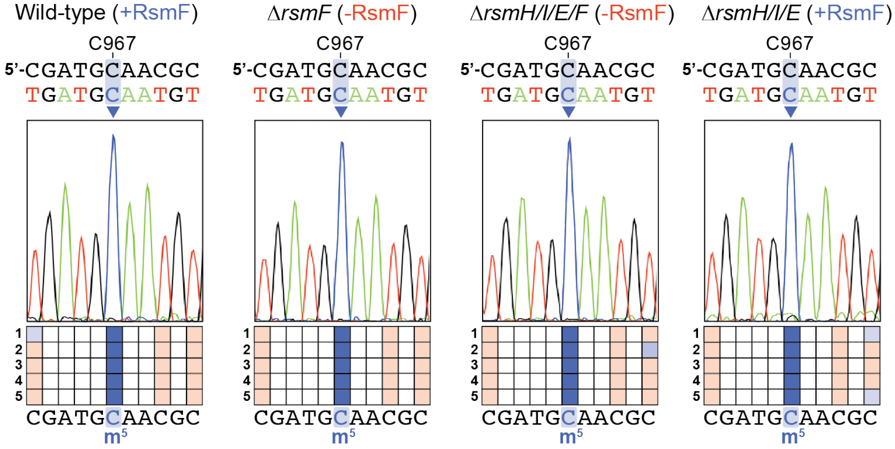


**Fig. S2. C967 methylation is unperturbed by the h44 16S rRNA methyltransferase gene deletions.** RNA bisulfite sequencing analysis of the 16S rRNA region surrounding nucleotide C967 (modified to $m^5C$ by RsmB which is present in all strains), shown for (**A**) wild-type 30S, (**B**) Δ*rsmF* 30S, (**C**) Δ*rsmH/I/E/F* 30S, and (**D**) Δ*rsmH/I/E* 30S. In each panel, the 16S rRNA sequence is shown (*top*) with the sequencing color-coded bisulfite converted sequence below. Also shown are representative sequencing chromatograms (*middle*) and grid maps summarizing C to T conversion at each cytosine position in the individual sequenced clones 1-5 (*bottom*). Converted sites (C to T) are shown in red and unconverted sites in blue, with the C967 position indicated and highlighted with the dark blue shade.

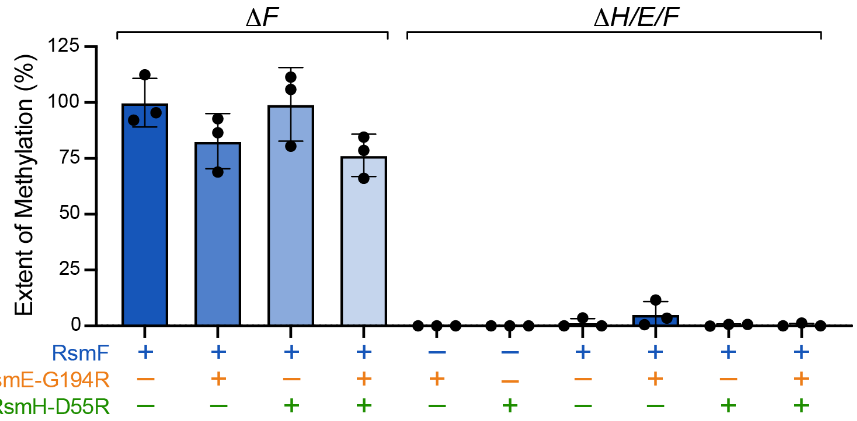


**Fig. S3. Catalytically inactive RsmE and RsmH enzymes do not rescue RsmF activity.** *In vitro* methyltransferase activity assays evaluating RsmF activity on 30S subunits isolated from Δ*rsmF* and Δ*rsmH/E/F* strains either with (+) or without (–) the addition of catalytically inactive enzyme variants, RsmE-G194R and RsmH-D55R, alongside controls without (–) the addition of active RsmF demonstrating that these variant enzymes are catalytically inactive. Bar graphs represent the mean percentage of methylation relative to the optimal Δ*rsmF* substrate.

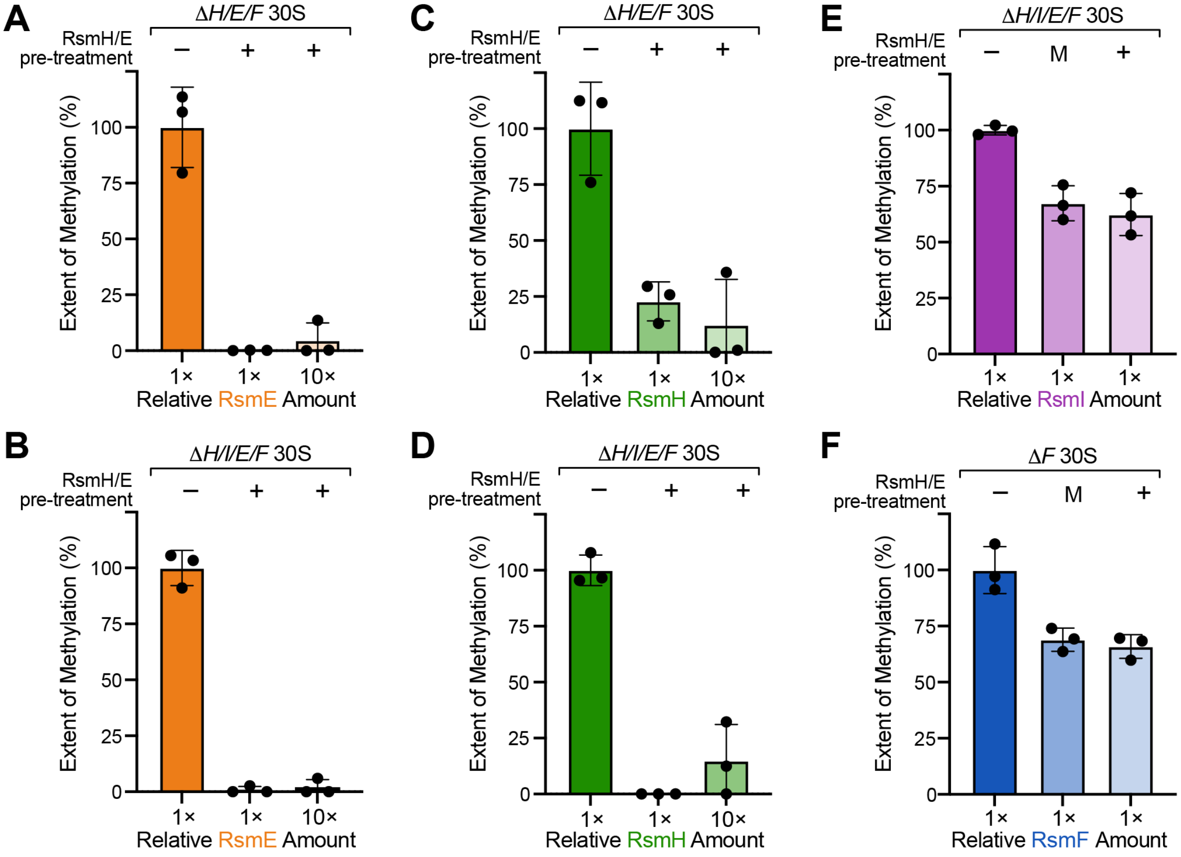


**Fig. S4. Decrease in baseline activity after RsmH and RsmE pre-treatment of 30S ribosomal subunits.** *In vitro* methyltransferase activity assays evaluating (**A**–**D**) completion of pre-treatment methylation (using non-radioactive SAM) for (**A**) RsmE on Δ*rsmH/E/F* 30S, (**B**) RsmE on Δ*rsmH/I/E/F* 30S, (**C**) RsmH on Δ*rsmH/E/F* 30S and (**D**) RsmH on Δ*rsmH/I/E/F* 30S, either untreated (–) or pre-treated (+) with RsmH and RsmE. Assays were conducted using either standard (1×) or a ten-fold higher (10×) relative concentration of enzyme. (**E**–**F**) Assessment of pre-treatment impact on substrate integrity for (**E**) RsmI on Δ*rsmH/E/F* 30S and (**F**) RsmF on Δ*rsmF* 30S, evaluated on untreated (–), mock-treated (M), and pre-treated (+) substrates. Activity is expressed as a percentage relative to the untreated substrate.

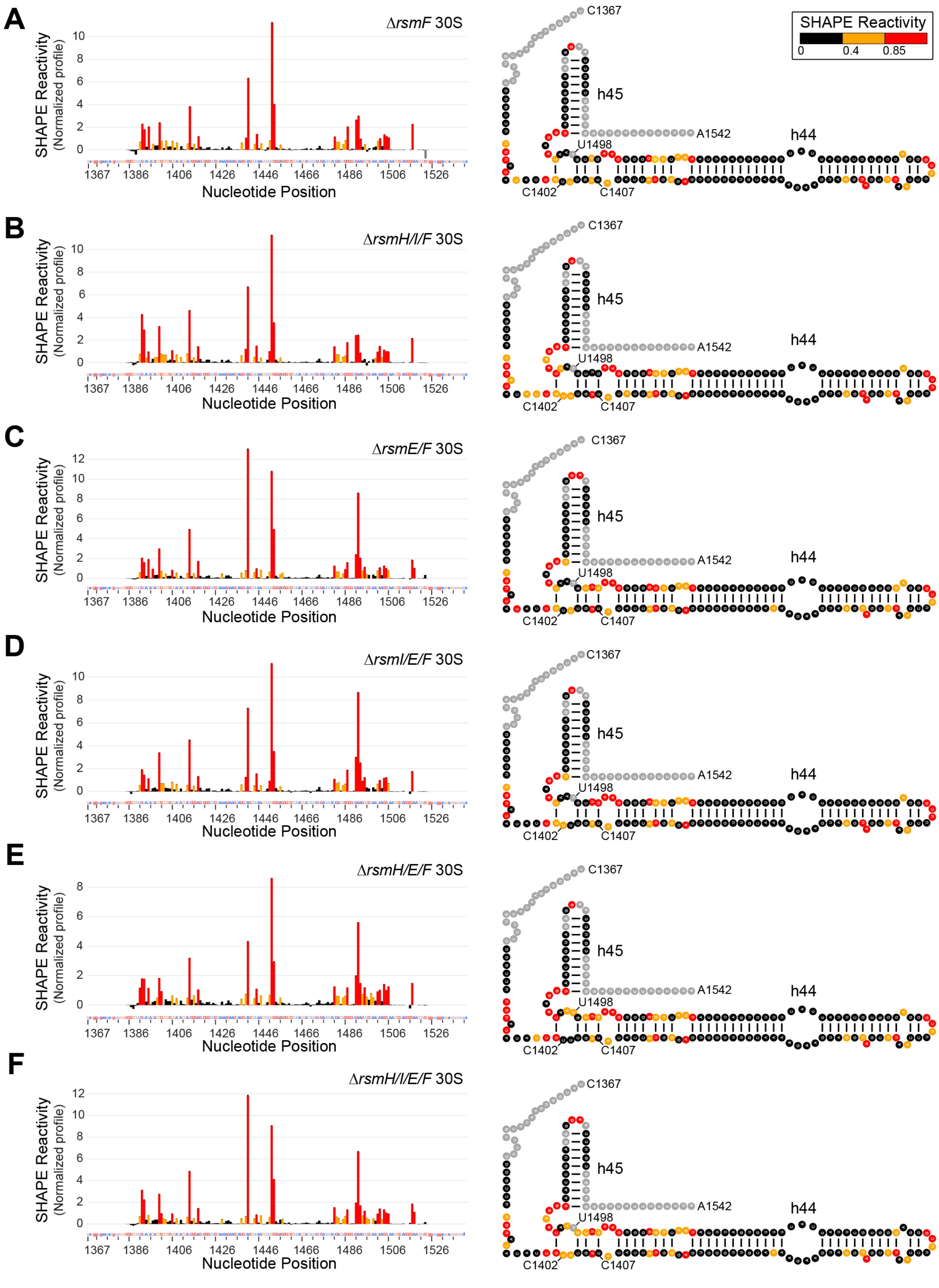


**Fig. S5. Normalized and averaged SHAPE reactivities of hypomethylated 30S subunits.** *In vitro* SHAPE-MaP analysis showing the reactivity profile of the 16S rRNA region encompassing h44 and h45 (nucleotides 1367–1542) corresponding to (**A**) Δ*rsmF* 30S, (**B**) Δ*rsmH/I/F* 30S, (**C**) Δ*rsmE/F* 30S, (**D**) Δ*rsmI/E/F* 30S, (**E**) Δ*rsmH/E/F* 30S, and (**F**) Δ*rsmH/I/E/F* 30S. Left panels display the normalized SHAPE reactivity profile across the targeted nucleotide positions. Right panels show the absolute reactivities mapped onto the corresponding 16S rRNA secondary structure. Nucleotides are color-coded based on their reactivity levels: low or no reactivity (0 to < 0.4; black), intermediate reactivity (0.4 to < 0.85; yellow), and high reactivity (0.85 to > 1.0; red). All profiles represent the average n = 3 biological replicates, with the exception of Δ*rsmI/E/F* strain (n=2).

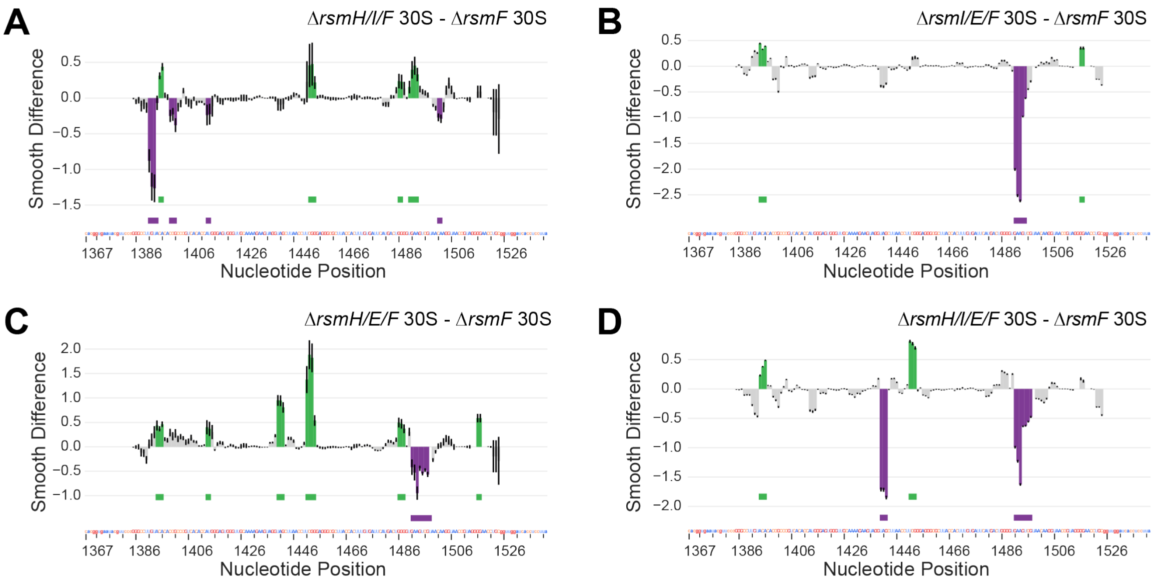


**Fig. S6. deltaSHAPE analysis results for hypomethylated 30S subunits.** deltaSHAPE profiles show the smoothed differences in nucleotide reactivity 16S rRNA region encompassing helices 44 and 45 (nucleotides 1367–1542). Profiles represent the differential reactivity of (**A**) Δ*rsmH/I/F*, (**B**) Δ*rsmI/E/F*, (**C**) Δ*rsmH/E/F*, and (**D**) Δ*rsmH/I/E/F* relative to Δ*rsmF*. Green indicates low differential reactivity, while purple indicates high differential reactivity. [Note that the Δ*rsmE/F* 30S analysis is excluded here as no reactivity differences meeting the 1.0 smoothing threshold were identified. Also see **Fig. S7** for analysis at the lower threshold of 0.725.].

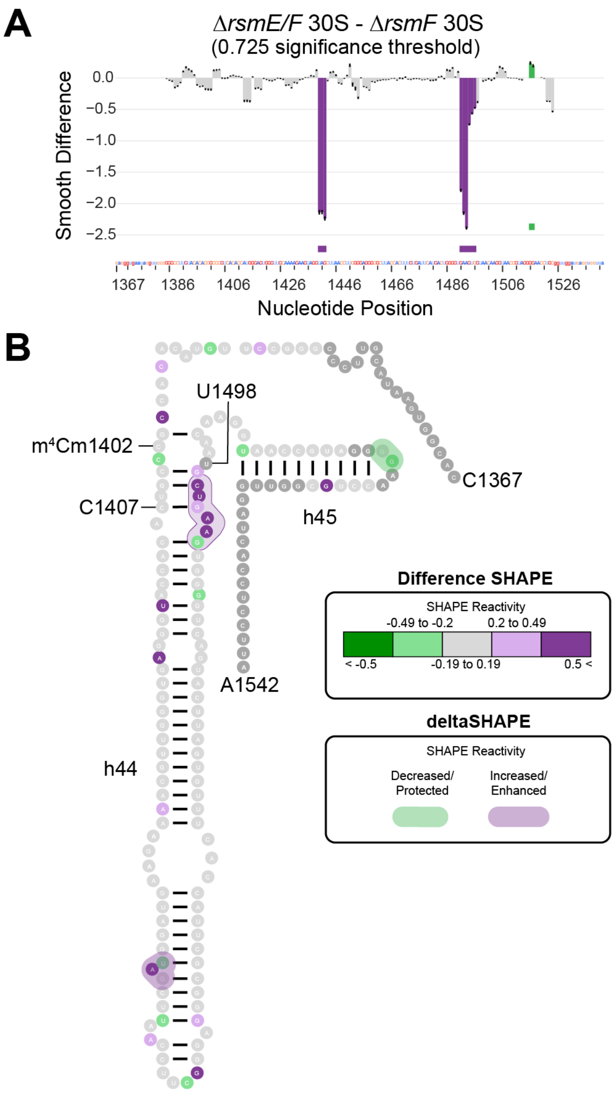


**Fig. S7. Nucleotide dynamics of the hypomethylated Δ*rsmE/F* 30S subunit.** *In vitro* SHAPE-MaP reactivity mapping of the 16S rRNA region encompassing h44 and h45 (nucleotides 1367–1542 representing the differential reactivity of the Δ*rsmE/F* mutant relative to Δ*rsmF*. (**A**) deltaSHAPE profile showing the smoothed differences in nucleotide reactivity. (**B**) Subtracted SHAPE reactivity projected onto the secondary structure. Differences are represented using both Difference SHAPE (individual circle colors) and deltaSHAPE (continuous structural shading over the circles). Green indicates a decrease in differential SHAPE reactivity, while purple indicates an increase in differential SHAPE reactivity. Data was analyzed at a 0.725 significance threshold.

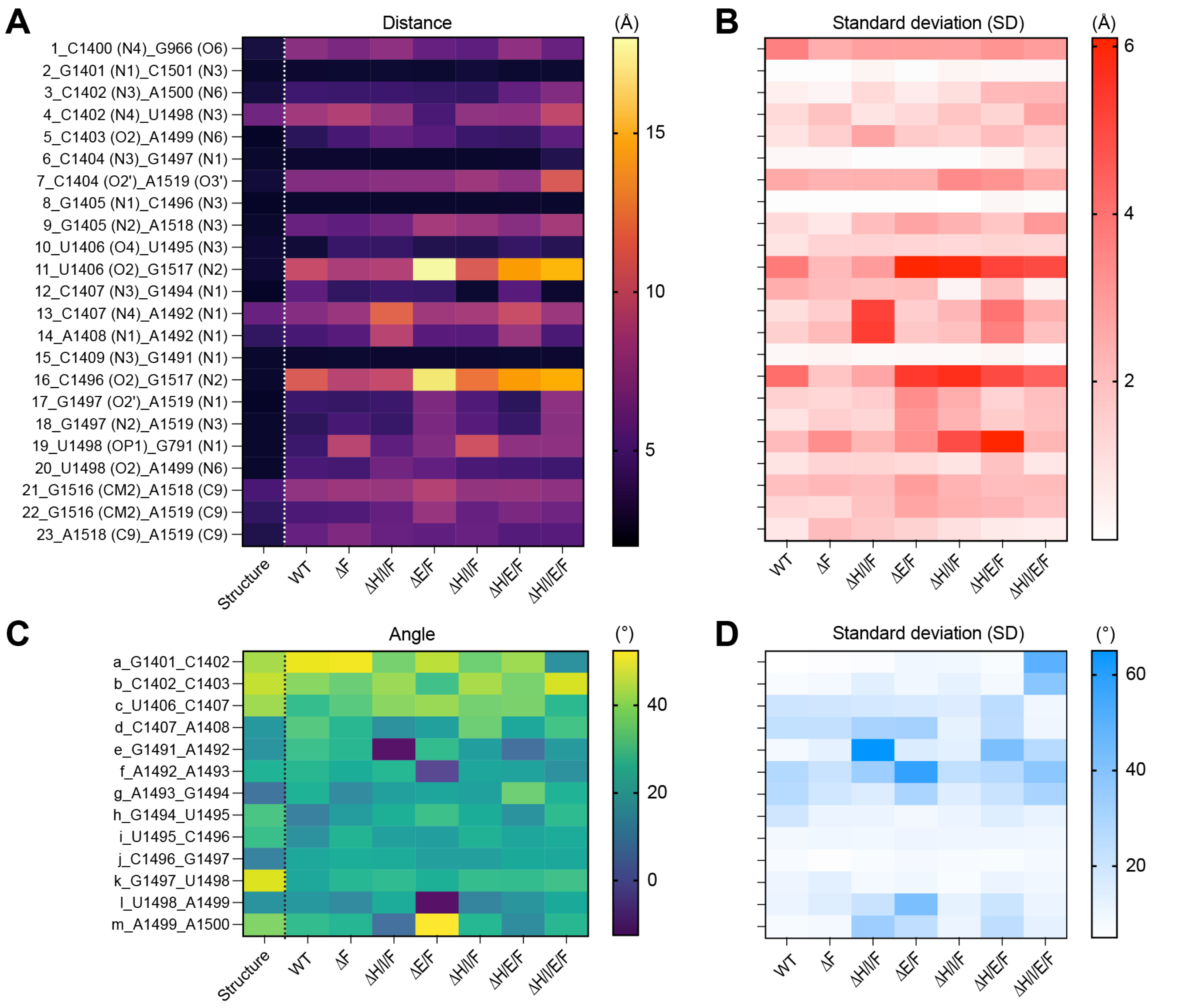


**Fig. S8. Distance and pseudodihedral angle measurements from MD simulation of hypomethylated 30S subunits.** Heat map representations of the measured parameters derived from the seven simulated systems with distinct h44 methylation status (indicated by the deletion strain) plus the starting structure before simulation (PDB 9Q87; denoted "Structure"): (**A**) average interatomic distances (Å) and (**B**) their associated standard deviations; (**C**) average pseudodihedral angles (˚) and (**D**) their standard deviations. Numbering and lettering of the distances and angles, respectively, correspond to the schematic in **Fig. 6A**; data shown for Δ*F* and Δ*HEF* are the same as in **Fig. 6B,C**.

### Supplementary Tables

**Table S1. List of primers used in study**

| Primer | DNA Sequence (5' to 3') | Description |
|---|---|---|
| | *Genotype Validation* | |
| *rsmH*-For | GACTTATCGGAGCGACTGC | Anneal to outside the ORF of gene to verify both replacement of gene and successful flipping of kanR cassette |
| *rsmH*-Rev | GAGCTTCTGTCACTCTGCTG | |
| *rsmE*-For | CTACACTAGCGGGATTCTTTTTG | |
| *rsmE*-Rev | CTGGAATCTTTCTTGATGTTGATG | |
| *rsmF*-For | CCAAATAATGCCCACTGCTCCG | |
| *rsmF*-Rev | CCGATCGCACCATTTTTCAGCC | |
| | *Rsm Gene Cloning* | |
| RsmI-ORF-For | GAAGATCTCAT**ATG**AAACAACACCAATCGGC | Clone *rsmI* into pET-44 and verify transfer of *rsmI*-Cam$^R$ genotype. **Start**/ **End** of *rsmI* ORF. |
| RsmI-ORF-Rev | CTAAAGCTTTTA**TTA**CCCCTGCTGCTCC | |
| RsmH-ORF-For | CGTGGATCCCATATG**ATG**GAAAACTATAAACATAC TAC | Clone *rsmH* into pET-44. **Start**/ **End** of *rsmH* ORF. |
| RsmH-ORF-Rev | CTCAAGCTTTTA**TTA**TGCATTCGTCCTCTCTG | |
| RsmE-ORF-For | CAAGGATCCCAT**ATG**CGTATCCCCC | Clone *rsmE* into pET-44. **Start**/ **End** of *rsmE* ORF |
| RsmE-ORF-Rev | ATCGGCCGTTA**TTA**GCCCAAATCGC | |
| RsmF-ORF-For | TAAGGATCCCAT**ATG**GCCCAACACACC | Clone *rsmF* into pET-44. **Start**/ **End** of *rsmF* ORF |
| RsmF-ORF-Rev | CTCAAGCTTTTA**TTA**GGCGTTACCGGTAAAAAGTT TC | |
| | *Mutagenesis (MEGAWHOP)* | |
| pET-44-For | GCAACCGCGAATTCACCTGTGG | Universal forward primer |
| G194R-RsmE-Rev | CCTTCCGG**ACG**AATCAGCAG | Reverse primer for first-round PCR in MEGAWHOP contains the altered codon (bold/ underlined) |
| D55R-RsmH-Rev | GGTCGCG**ACG**GATCGCCAG | |
| | *16S rRNA Bisulfite Sequencing* | |
| C967 Region-RT/Rev | CTCTAAAAACTTCCATAA | RT bisulfite-treated 16S rRNA (~100 nt region encompassing C967) and PCR of cDNA |
| C967 Region-For | ATGAATTGATGGGGGTTT | |
| C1407 Region-RT/Rev | TAAACACCCTCCCAAAAA | RT bisulfite-treated 16S rRNA (~100 nt region encompassing C1407) and PCR of cDNA |
| C1407 Region-For | TGAATATGTTTTTGGGTT | |
| | *SHAPE-MaP* | |
| RT Primer | TAAGGAGGTGATCCAACC | Reverse transcription of modified 16S rRNA |
| Δ*rsmF*_2A3-For | *TCGTCGGCAGCGTCAGATGTGTATAAGAGACAG* **AGGCAGAA**CACGGTGAATACGTTCCC | Amplification of cDNA resulting from 2A3 or DMSO-treated reverse transcribed 16S rRNA, with Nextera adapter (*italicized*) and inline barcode (**bold**) sequences. |
| Δ*rsmF*_2A3-Rev | *GTCTCGTGGGCTCGGAGATGTGTATAAGAGACAG* **GTAAGGAG**TAAGGAGGTGATCCAACC | |
| Δ*rsmF*_DMSO-For | *TCGTCGGCAGCGTCAGATGTGTATAAGAGACAG* **TAGGCATG**CACGGTGAATACGTTCCC | |
| Δ*rsmF*_DMSO-Rev | *GTCTCGTGGGCTCGGAGATGTGTATAAGAGACAG* **CTAAGCCT**TAAGGAGGTGATCCAACC | |
| Δ*rsmH/I/F*_2A3-For | *TCGTCGGCAGCGTCAGATGTGTATAAGAGACAG* **GGACTCCT**CACGGTGAATACGTTCCC | |
| Δ*rsmH/I/F*_2A3-Rev | *GTCTCGTGGGCTCGGAGATGTGTATAAGAGACAG* **AAGGAGTA**TAAGGAGGTGATCCAACC | |
| Δ*rsmH/I/F*_DMSO-For | *TCGTCGGCAGCGTCAGATGTGTATAAGAGACAG* **TAGGCATG**CACGGTGAATACGTTCCC | |
| Δ*rsmH/I/F*_DMSO-Rev | *GTCTCGTGGGCTCGGAGATGTGTATAAGAGACAG* **CTAAGCCT**TAAGGAGGTGATCCAACC | |

| | |
|---|---|
| Δ*rsmE/F*_2A3-For | *TCGTCGGCAGCGTCAGATGTGTATAAGAGACAG* **GGAGCTAC**CACGGTGAATACGTTCCC |
| Δ*rsmE/F*_2A3-Rev | *GTCTCGTGGGCTCGGAGATGTGTATAAGAGACAG* **TTCTAGCT**TAAGGAGGTGATCCAACC |
| Δ*rsmE/F*_DMSO-For | *TCGTCGGCAGCGTCAGATGTGTATAAGAGACAG* **ACTCGCTA**CACGGTGAATACGTTCCC |
| Δ*rsmE/F*_DMSO-Rev | *GTCTCGTGGGCTCGGAGATGTGTATAAGAGACAG* **TCGACTAG**TAAGGAGGTGATCCAACC |
| Δ*rsmI/E/F*_2A3-For | *TCGTCGGCAGCGTCAGATGTGTATAAGAGACAG* **AGGCAGAA**CACGGTGAATACGTTCCC |
| Δ*rsmI/E/F*_2A3-Rev | *GTCTCGTGGGCTCGGAGATGTGTATAAGAGACAG* **GTAAGGAG**TAAGGAGGTGATCCAACC |
| Δ*rsmI/E/F*_DMSO-For | *TCGTCGGCAGCGTCAGATGTGTATAAGAGACAG* **TCCTGAGC**CACGGTGAATACGTTCCC |
| Δ*rsmI/E/F*_DMSO-Rev | *GTCTCGTGGGCTCGGAGATGTGTATAAGAGACAG* **ACTGCATA**TAAGGAGGTGATCCAACC |
| Δ*rsmH/E/F*_2A3-For | *TCGTCGGCAGCGTCAGATGTGTATAAGAGACAG* **CGTACTAG**CACGGTGAATACGTTCCC |
| Δ*rsmH/E/F*_2A3-Rev | *GTCTCGTGGGCTCGGAGATGTGTATAAGAGACAG* **TATCCTCT**TAAGGAGGTGATCCAACC |
| Δ*rsmH/E/F*_DMSO-For | *TCGTCGGCAGCGTCAGATGTGTATAAGAGACAG* **GGACTCCT**CACGGTGAATACGTTCCC |
| Δ*rsmH/E/F*_DMSO-Rev | *GTCTCGTGGGCTCGGAGATGTGTATAAGAGACAG* **AAGGAGTA**TAAGGAGGTGATCCAACC |
| Δ*rsmH/I/E/F*_2A3-For | *TCGTCGGCAGCGTCAGATGTGTATAAGAGACAG* **CGGAGCCT**CACGGTGAATACGTTCCC |
| Δ*rsmH/I/E/F*_2A3-Rev | *GTCTCGTGGGCTCGGAGATGTGTATAAGAGACAG* **GCGTAAGA**TAAGGAGGTGATCCAACC |
| Δ*rsmH/I/E/F*_DMSO-For | *TCGTCGGCAGCGTCAGATGTGTATAAGAGACAG* **GCGTAGTA**CACGGTGAATACGTTCCC |
| Δ*rsmH/I/E/F*_DMSO-Rev | *GTCTCGTGGGCTCGGAGATGTGTATAAGAGACAG* **CCTAGAGT**TAAGGAGGTGATCCAACC |

**Table S2. List of buffers used in study**

| Buffer Name | pH | Composition |
|---|---|---|
| Cell Lysis Buffer (RsmH/I/E/F) | 7.6 | 50 mM HEPES-NaOH, 1 M NaCl, 10 mM imidazole, 0.5% Triton X-100, 2 mM $\beta$-mercaptoethanol |
| High Salt Dialysis Buffer (RsmH/I/E/F) | 7.6 | 50 mM HEPES-NaOH, 2 M NaCl, 10 mM imidazole, 2 mM $\beta$-mercaptoethanol |
| HisTrap Buffer A (RsmH/I/E/F) | 7.6 | 50 mM HEPES-NaOH, 500 mM NaCl, 10 mM imidazole, 2 mM $\beta$-mercaptoethanol |
| HisTrap Buffer B (RsmH/I/E/F) | 7.6 | 50 mM HEPES-NaOH, 500 mM NaCl, 1 M imidazole, 2 mM $\beta$-mercaptoethanol |
| Gel Filtration (GF) Buffer (Rsm E/F) | 7.6 | 20 mM HEPES-NaOH, 150 mM NaCl, 2 mM $\beta$-mercaptoethanol |
| Gel Filtration (GF) Buffer (RsmH/I) | 7.6 | 10 mM HEPES-NaOH, 300 mM NaCl, 2 mM $\beta$-mercaptoethanol |
| Buffer 1 (30S Purification) | 7.6 | 10 mM HEPES, 1 M $NH_4Cl$, 10 mM $MgCl_2$, 6 mM $\beta$-mercaptoethanol |
| Buffer 2 (30S Purification) | 7.6 | 10 mM HEPES, 100 mM $NH_4Cl$, 10 mM $MgCl_2$, 6 mM $\beta$-mercaptoethanol |
| Buffer 3 (30S Purification) | 7.62 | 10 mM HEPES, 100 mM $NH_4Cl$, 0.3 mM $MgCl_2$, 6 mM $\beta$-mercaptoethanol |
| Buffer G | 7.5 | 5 mM HEPES-KOH, 50 mM KCl, 10 mM $NH_4Cl$, 10 mM MgOAc, 6 mM $\beta$-mercaptoethanol |
| MaP Buffer | 8 | 50 mM Tris-HCl, 75 mM KCl, 6 mM $MnCl_2$, 10 mM DTT, 0.5 mM dNTPs. |
